# Lung-mimicking 3-Dimensional hydrogel culture system recapitulates key tuberculosis phenotypes and demonstrates pyrazinamide efficacy

**DOI:** 10.1101/2023.01.24.525291

**Authors:** Vishal K. Gupta, P.S. Abhirami, Vaishnavi V. Vijaya, K.M. Jyothsna, Sharumathi Jeyasankar, Varun Raghunathan, Rachit Agarwal

**Author notes:** Corresponding author (Rachit Agarwal).

## Abstract

Faithful mimics of tuberculosis (TB) infection are needed to provide mechanistic insights into the complex host-pathogen interactions and accelerate drug discovery. Current *in vitro* models only allow short investigation durations, present divergent transcriptional signatures to human infections, and are unreliable drug discovery platforms. We developed a 3D collagen culture system mimicking the lung microenvironment (collagen fibres, pore size and stiffness), where we incorporated *Mycobacterium tuberculosis* (Mtb) infected human THP-1 or primary monocytes. Dual RNA-sequencing revealed high mammalian gene expression similarity with patient samples compared to 2D macrophage infections. Similarly, gene expression of bacteria was much more representative to *in vivo* gene expression compared to bacteria in 2D cultures (114 genes in 3D vs 21 genes in 2D). Key phenotypes observed in humans, such as foamy macrophages and mycobacterial cords (never seen in any other *in vitro* culture system), were reproduced in our model. Our system overcomes many challenges associated with the traditional platforms, including showing remarkable efficacy with clinically relevant concentrations of first-line anti-TB drug pyrazinamide, not seen in any other *in vitro* model, making it reliable, readily adoptable for tuberculosis studies and drug screening.

**Significance statement:** Mtb is a slow-growing pathogen which modulates host response over time. The current *in vitro* platforms offer a very short study duration to study, are unreliable as drug discovery platforms, and the phenotypic and genotypic traits of the host and pathogen differ. The collagen-I hydrogel culture system developed in this study addresses these challenges by successfully recapitulating several key phenotypes observed in human infections. Dual RNA sequence also showed excellent gene expression similarities for both the host and the bacteria. Furthermore, remarkable efficacy with the antibiotic Pyrazinamide was demonstrated, a first for *in vitro* cultures despite over 50 years of clinical use of the drug. We expect our platform to be exploited widely for drug discovery and understanding host-pathogen interactions.

## Introduction

In 2021, the World Health Organization (WHO) reported 10 million new TB cases, which resulted in 1.5 million deaths (1). TB is transmitted through aerosols containing the bacteria generated by an infected individual. Upon reaching the alveolus of the new host, the bacteria infect the resident alveolar macrophages, which migrate to the lung interstitium and recruit circulating monocytes and other immune cells, leading to the formation of granuloma (2, 3). Monocytes and macrophages carry a large reservoir of bacteria *in vivo* and play critical roles in the host response to mycobacterial infection (4–6). These cells are important for studying host-pathogen interactions and hence have been widely employed in the field (7–9).

Conventionally*, in vitro* Mtb infection experiments are performed in two-dimensional (2D) tissue culture systems on polystyrene plates, using primary human monocyte-derived macrophages, mouse bone marrow-derived macrophages (BMDMs), and human THP-1 macrophages (10). These studies have contributed to our understanding of complex processes such as the arrest of phagosomal maturation, activation of innate immune responses, and the underlying mechanisms modulating various cell death pathways (7). However, the existing culture systems have critical drawbacks: First, monolayer (2D) cultures fail to provide the natural environments for the cells, wherein cells are usually subjected to very stiff surfaces (∼10 GPa), resulting in many dysregulated cellular processes (11). Second, host cell viability after Mtb infection is low in 2D cultures, limiting the length of the experiments to 4-6 days. Mtb is a slow-growing pathogen (median doubling time of >100 h) (12) that modulates host responses to help its survival over several days/weeks. The conventional 2D platforms fail to recapitulate these intricate host-pathogen dynamics. Third, substantial variation exists in gene expression between infected cells in 2D cultures and human infection (13). Fourth, the current systems seem unreliable as platforms to screen TB drugs. For instance, pyrazinamide, a first-line anti TB drug, has shown little to no efficacy in any previously used *in vitro* culture system at clinically relevant concentrations but is extremely effective *in vivo* (14). Similarly, several clinical trials with potential TB drugs and treatment regimens have failed despite showing promise in the initial stage of testing (15). These limitations underscore the need to develop better and more representative culture systems to accurately study host-mycobacteria interactions. An ideal culture system should (a) provide a more extended timeframe (∼weeks) for studying host-pathogen dynamics in TB; (b) be able to better recapitulate the genotypic and phenotypic characteristics of human infection; (c) be a more reliable platform for advancing drug susceptibility studies; and (d) be simple with accessible components to enable easy and widespread adoption in the field.

In recent years, three-dimensional (3D) *in vitro* models have gained significant interest in the field and can act as an alternative to animal testing for initiating clinical trials as per the FDA Modernization Act 2.0 (16, 17). 3D models provide a more realistic representation of human tissues than their 2D counterparts, as they mimic cells *in vivo* that are surrounded by other cells and the extracellular matrix (ECM) (18). A major advancement in using ECM-based matrices to study TB host-pathogen interaction was achieved by Kapoor *et al.* (19), where the authors used collagen scaffolds to infect peripheral blood mononuclear cells (PBMCs) with Mtb leading to the formation of cellular aggregates resembling TB granulomas. The authors demonstrated that their model could recapitulate the dormant bacterial phenotype. However, biomechanical aspects, gene expression comparison with primary human samples and pyrazinamide efficacy were not reported. Another important study used spherical alginate gels doped with collagen as a 3D matrix to infect PBMCs with Mtb to model TB granuloma (19, 20). However, since alginate is not part of the lung ECM, it could result in non-physiological signalling. The synthesis of alginate spheres also requires access to specialized equipment that may not be readily available in BSL-3 settings. This latter study also showed the efficacy of first-line anti-TB drugs in their model including pyrazinamide. However, the concentrations required to reduce bacterial viability were much higher (500 µg/mL) than the clinically relevant concentrations of the drug (50 µg/mL). We aimed to engineer a readily adoptable collagen-based culture system that mimics human TB infection and recapitulates gene expression, bacterial and mammalian phenotypes and drug efficacy.

We have synthesized hydrogels made of Collagen I, the primary extracellular component in the human lungs, and tuned the micromechanical environment similar to human lung tissue. The collagen gels are easy to form and do not require any specialized equipment. These gels supported the culture of THP-1 and primary human monocytes, with the cells sustaining Mtb infection beyond 3 weeks. Dual RNA-sequencing of the host and pathogen transcriptomes from collagen gels showed that the gene expression profiles of the mammalian and bacterial cells infected in our collagen gels mimic human infection significantly better than in 2D cultures. The monocytes in the infected gels differentiated into macrophages and accumulated lipid bodies, a phenotype observed in macrophages in human TB granulomas (21). The longevity of this culture system enabled us to see bacterial growth in the form of cords, a phenotype observed *in vivo* (5, 22-24) but not in any *in vitro* macrophage infection model. Finally, our culture system emerged as an excellent drug testing platform, shown by the efficacy of pyrazinamide, at or below the concentrations used clinically (50 µg/mL). Rapid and widespread adoption of this system is thus likely and would not only aid drug discovery and development but also shed new insights into the complex host-pathogen interactions underlying TB infections.

## Results

### 1. Characterization of collagen gels revealed their micromechanical similarity with human lungs and cytocompatibility

Mtb infection manifests in the lung interstitium (3), primarily comprised of the ECM network of collagen I and III (25, 26) and other proteoglycans. We synthesized and tuned a physically crosslinked collagen I hydrogel that can mimic the biochemical and biophysical cues of human lung interstitium (27, 28). To characterize the mechanical microenvironment, collagen gels were analyzed using Atomic Force Microscopy (AFM). The Young’s modulus obtained was 0.64 kPa ± 0.12 kPa and 2.1 kPa ± 0.3 kPa for 1 mg/mL and 2 mg/mL collagen gels, respectively (**Fig. 1A**). Since the modulus of healthy human lung tissue is in range of 1.4 to 7 kPa (29, 30), we chose 2 mg/mL collagen gels for all subsequent assays. Next, to characterize whether these gels can mimic the fibrillar structure of ECM in human lungs, the collagen gels were imaged using second harmonic generation (SHG) imaging, a label-free imaging technique based on collagen’s native property to generate the second harmonics when when incident with a high intensity laser beam (31). SHG imaging showed distinct fibrils in collagen hydrogels (**Fig. 1B**) as has been widely reported (27, 32) and morphologically mimicked lung ECM microenvironment. The average pore area in gels with final collagen I concentration of 2 mg/mL was 8.45 µm^2^ ± 2.37 µm^2^, allowing cells to move through the matrix.

**Fig. 1.**
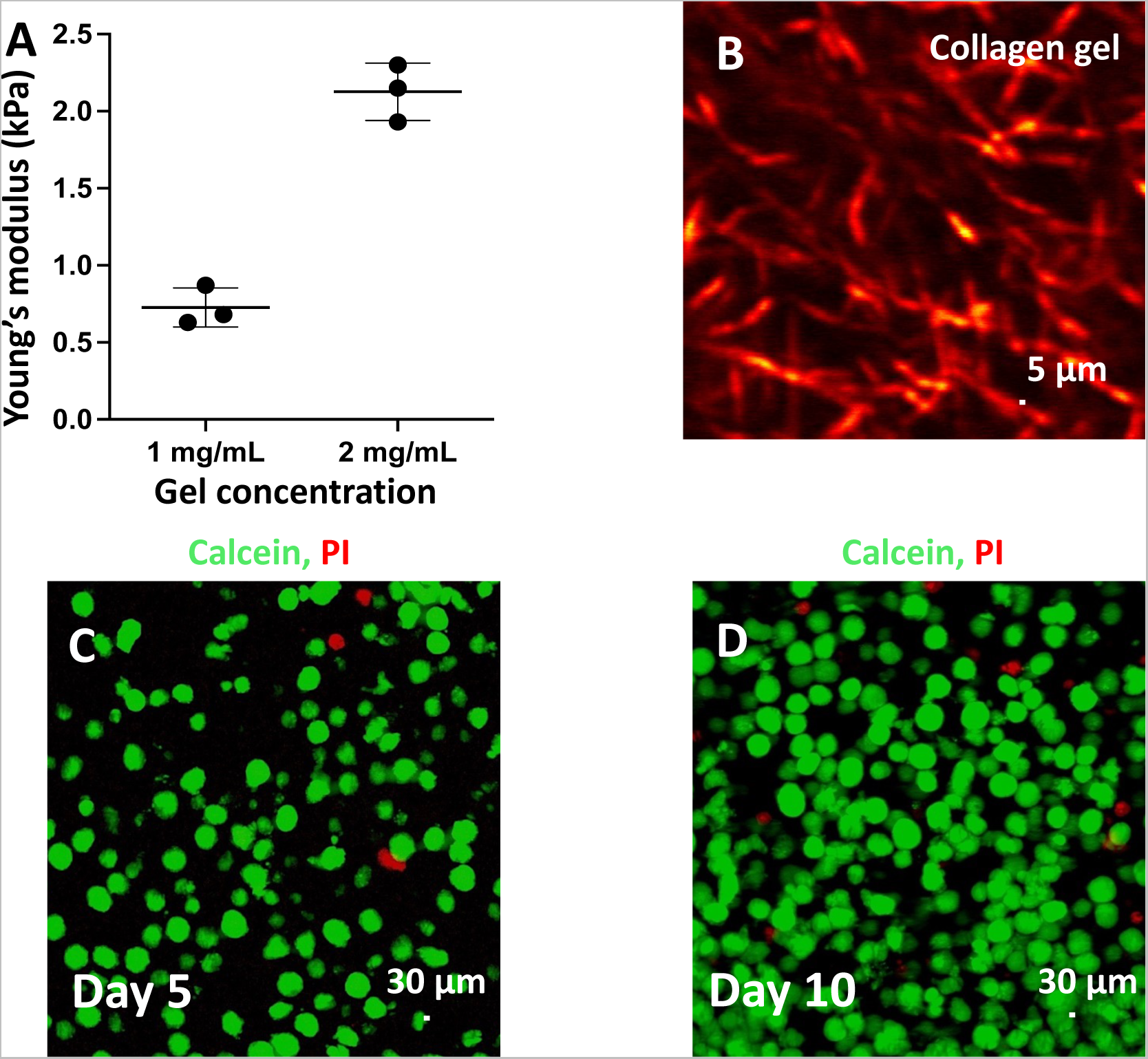
Collagen gels mimic the micromechanical environment of lungs and are cytocompatible. (**A**) Young’s modulus of 1 mg/mL and 2 mg/mL collagen gels. Data in graphs represent the mean ± s.d. (*n* = 3). (**B**) Representative SHG image of 2 mg/mL collagen gel. Scale: 5 µm. (**C**) and (**D**) Representative images of Calcein-AM (green, live) – PI (red, dead) stained cells images in collagen gels on days 5 and 10, respectively.

The ECM is critical to cell viability as cellular detachment from ECM alters their processes, such as metabolism. The separation from ECM also leads to cell death due to anoikis (33). Collagen plays a critical role in the progress of TB infection as its degradation results in reduced cell viability leading to the caseation of TB granuloma (34). We first tested the cytocompatibility of collagen gels for THP-1 cells by culturing them for 10 days. Minimal cytotoxicity was observed by Calcein -AM – propidium iodide (PI) staining (**Fig. 1C and 1D**, Day 5 – 92.3 ± 2.76 % viability, Day 10 – 87.1± 4.37 viability).

### 2. Collagen gels support monocyte infection and cell survival for a prolonged duration

We characterized the infection percentages at a range of MOIs (0.1- 40) in PMA-treated THP- 1 macrophages (2D infection) and for our 3D model. To achieve infection in 3D, THP-1 monocytes and Mtb were added to the collagen gel before polymerization. We used monocytes to mimic the natural process of TB infection *in vivo*, during which inflammation leads to the recruitment of circulating monocytes to the site of infection (35). Successful infection was achieved upon incubating THP-1 cells with Mtb in the hydrogels for 12 h, after which the gels were treated with a mammalian cell impermeable antibiotic (amikacin sulfate) to eliminate extracellular bacteria (36) (**Fig. 2A**, **Supplementary Video 1 and 2**). An MOI of 40 in the 3D model resulted in a similar intracellular infection percentage as an MOI of 1 in 2D infection (**Supplementary Fig. S1**) and was used for further experiments. Harmonizing the initial infection percentages in 2D and 3D was done for allowing subsequent comparisons. We also found that infecting undifferentiated monocytes with Mtb (MOI of 40) in the absence of collagen gels resulted in very few THP-1 cells getting infected (**Supplementary Fig. S2)**, which highlights the importance of collagen in mediating host-Mtb infection.

**Fig. 2.**
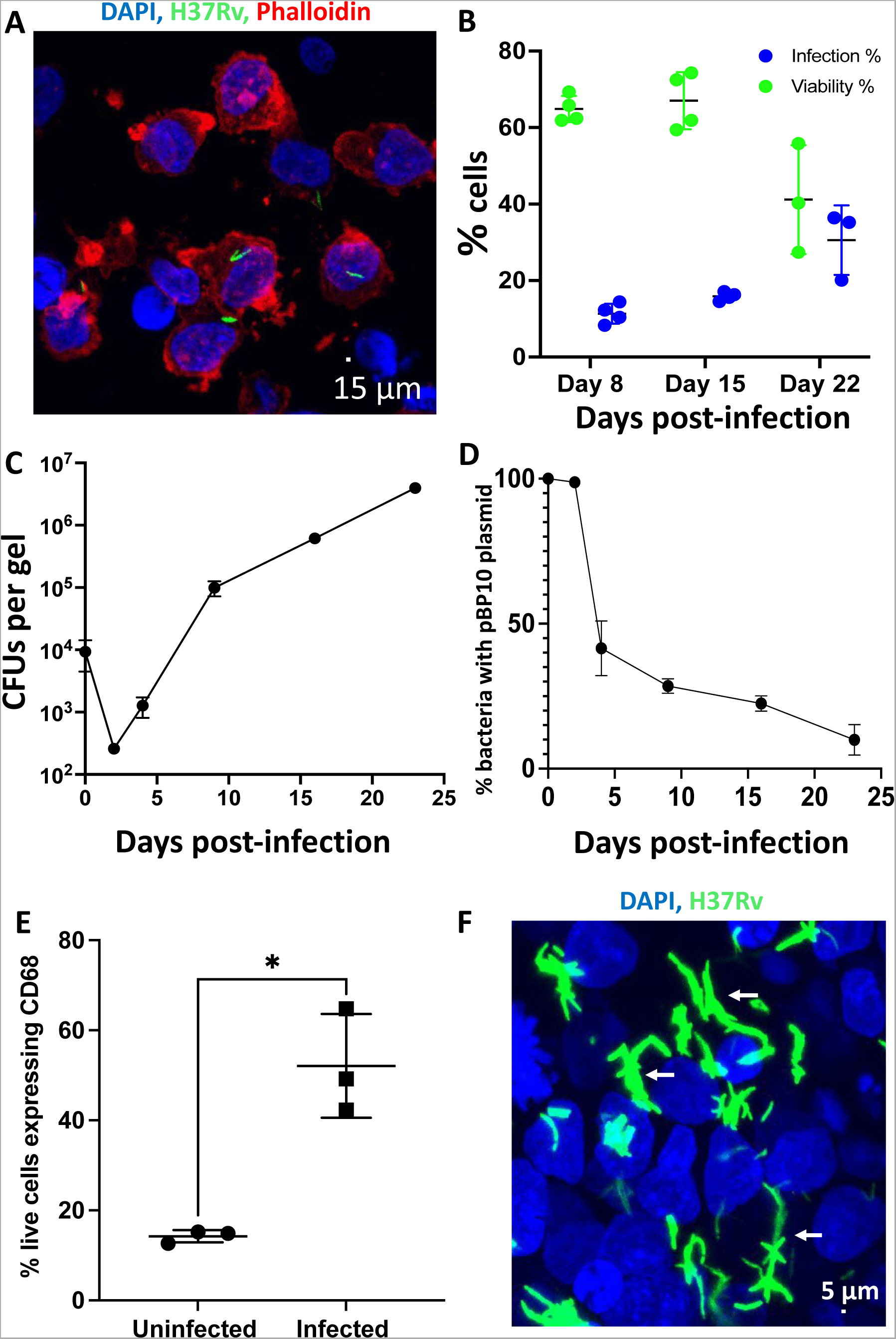
Collagen gels support THP-1 cell infection with Mtb over a long duration. (**A**) Representative fluorescence microscopy image of infected THP-1 cells in collagen gel (Hoechst-Blue), GFP-Mtb (H37Rv) (Green), and Phalloidin (Red) at 5 days post-infection. (**B**) The percentage of live THP-1 cells (green), and THP-1 cells infected with H37Rv-GFP (blue) at various intervals (*n* = 4) as determined using flow-cytometry assays (Gating strategy: **Supplementary Fig. 15**). (**C**) Kinetics of bacterial growth (CFUs) in the 3D collagen culture system (*n* = 3). (**D**) The percentage of intracellular bacteria containing the replication clock plasmid (pBP10) at various time points in the infected collagen gels, represented by CFUs (mean ± s.d.) on kanamycin vs non-kanamycin plates (*n* = 3). (**E**) Plot showing the percentage of live CD68-positive cells in uninfected and infected gels (*n* = 3) at 10 days post-infection. (**F**) Representative fluorescence image showing that Mtb (green) forms complex cord-like structures in the infected collagen gels (highlighted using white arrows). Images were acquired from infected collagen 14 days post-infection. Data in the graphs represent the mean ± s.d., and the *p-value* was determined by a two-tailed unpaired t-test with Welch’s correction using GraphPad Prism software. *p*-value < 0.05 was considered significant. \**p*=0.0282

A major limitation of the existing 2D infection models is the relatively small time frame available for conducting experiments (37). An infection with MOI of 1 resulted in viable host cells for approximately 7 days, beyond which the cell viability dropped below 10% (**Supplementary Fig. S3)**. This is a major constraint for long-term mechanistic and drug studies as Mtb is a slow-growing pathogen with a median doubling time >100 h inside mammalian cells (12). Post-infection, the bacteria take 1-2 days to adapt to the mammalian system (38), which further reduces the available time for assays by nearly 25-30%.

In the infected collagen 3D environment, we could successfully culture cells for 22 days (**Fig. 2B**). At various time points, the viability and infection rates were analyzed using flow cytometry. 8 days post-infection, 60±3% of cells were viable, and the corresponding numbers for 15- and 22 days post-infection were 63±7% and 40±9%, respectively. After 8, 15, and 22 days, the infection rates were 8±0.9%, 10±1.3%, and 30±12.4%, respectively. We also performed a colony forming unit (CFU) assay using H37Rv transformed with the pBP10 clock plasmid (39) (replication clock plasmid) to understand the bacterial growth kinetics (**Fig. 2C & 2D)**. We found that during the first 2 days of infection, the CFUs reduce with minimal bacterial replication, as seen from no loss of the replication clock plasmid, which could be due to the time bacteria needs to adapt to the system. As time progresses, Mtb grows (with a corresponding loss in clock plasmid) in the THP-1 cells cultured in collagen gels, and hence the proportion of infected cells increases with time while the overall viability of mammalian cells decreases. Thus, this platform offers a significantly longer window for testing drugs and studying host-pathogen interaction during infection than 2D culture systems. A similar long time window to study host-pathogen dynamics was shown when PBMCs were infected with Mtb in alginate-collagen matrices (20).

During lung infection, monocytes get recruited to the site of inflammation and differentiate into macrophages (40, 41). In the collagen culture system, to determine whether monocytes were differentiating into macrophages upon Mtb exposure, cells were stained with CD68, a specific intracellular marker for macrophages (20, 42). Mtb infection resulted in CD68 expression in nearly 52% of cells 10 days post-infection (**Fig. 2E**) compared to only 14% in uninfected gels. We then analyzed the Mtb positive vs bystander cells in the infected conditions using flow cytometry to determine if all cells were uniformly differentiated into macrophages. Nearly all Mtb-infected cells were positive for CD68 (∼95%) (**Supplementary Fig. S4**), implying that infection resulted in the differentiation of monocytes to macrophages in collagen gels. We also observed that about 42% of the bystander (uninfected) population showed CD68 expression (**Supplementary Fig. S4A**), implying that the infected microenvironment promoted the differentiation of THP-1 monocytes to macrophages. These results were similar to an earlier study which showed differentiation of primary human monocytes to macrophages upon infection with Mtb in alginate-collagen matrices (20). We also looked at the macrophage activation marker, iNOS, which has been shown to play an important role in mice, macaques and human TB (43–46). As expected, we observed a significant upregulation of iNOS upon Mtb infection in differentiated macrophages **(Supplementary Fig. S4B)**.

Mtb can aggregate and form cord-like structures, known as cording, which is regularly observed *in vivo* (22, 23, 47, 48) and is associated with the bacteria’s virulence and immune escape mechanism. Cording correlates to the growth of Mtb in the extracellular milieu *in vivo* (48, 49) and cords are also observed in human infection (47, 50). However, the role of cords in modulating host dynamics during infection is still to be established. We also observed bacterial cording at later time points in the culture system (post 10 days) (**Fig. 2F**) which were mostly extracellular **(Supplementary Fig. S5)**. We classified these elongated bacterial structures as cords if the Ferret diameter was above 10 and the average found in the system was 15.28 ± 2.17 μm, 14 days post-infection. These cords grew longer as the duration of the infection progressed. One of the hypotheses suggests that cording enables the bacteria to prevent phagocytosis once they become extracellular upon killing the host cell (50, 51) and have also been shown to cause macrophage lysis upon contact (52). Previous *in vitro* studies have shown that human lymphatic endothelial cells (50) and alveolar epithelial cell line (A549) are conducive for cord formation by Mtb, 3-4 days post-infection. Furthermore, previous work has shown that macrophages form extracellular traps in response to Mtb cords to restrict the pathogen’s growth; however, in this model, the mammalian cells were infected with cords grown in 7H9 without tween (as opposed to single cell suspension of bacteria) (52, 53). Hence, this is the first *in vitro* macrophage infection culture system to show the development of Mtb cord phenotype.

### 3. Dual RNA sequencing revealed similarities in gene expression between mammalian cells of the collagen hydrogel system and human TB granulomas

We carried out dual RNA sequencing to dissect the host-pathogen interaction mechanisms by identifying their transcriptomic differences in the collagen gels 14 days post-infection. The differentially expressed mammalian, and Mtb genes are listed in **Supplementary Table 1**. This time point was chosen as it allowed enough time for the bacteria to adapt to the host. The differentially expressed gene (DEG) distribution for mammalian genes is shown in **Fig. 3A**. Gene expression patterns from our system were compared with the expression profiles of an infected human 2D macrophage cell line (GSE162729)/ primary cells (GSE174566) and with human infection (caseous lung granuloma (GSE20050) and lymph-node granuloma (GSE17443). The 3D collagen culture system had the highest number of similarly regulated genes with human caseous TB granuloma (21) (gene list: **Supplementary Table 2**) and with human lymph node granuloma (13) (gene list: **Supplementary Table 3**) compared with the other 2D infection models (**Fig. 3B**), which had several fold less similarly regulated genes. The top upregulated and downregulated DEGs common between the lymph node granuloma and the 3D collagen culture system showed similar regulation patterns (**Fig. 3C**). To better understand the functional relevance of such a gene expression pattern, we performed a REACTOME pathway gene set enrichment analysis for the differentially regulated genes in various models and found a significant overlap in both the upregulated and downregulated pathways between the human TB granulomas and 3D collagen culture system (**Supplementary Fig. S6**). The upregulated REACTOME pathways in the collagen model corresponded to ECM organization, signal transduction, while the downregulated ones were associated with protein metabolism, organelle biogenesis, and maintenance, among others. The transcriptomic response of the PBMCs to Mtb in a 3D alginate matrix supplemented with collagen had also shown many pathways to be similarly regulated with lymph node TB granuloma. However, the ECM remodelling pathways which are a significant characteristic of primary human TB granulomas did not get enriched in their system probably due to the use of alginate in the matrix (13). In the collagen culture system, though we only used collagen 1 as the ECM, the transcriptomic data showed significant upregulation of genes involved in the synthesis of other ECM proteins such as fibronectin (FN1) and laminin (LAMA2, LAMA4), pointing towards the remodelling of ECM during infection in our model. The 2D models, on the other hand, showed contradictory regulation of many pathways such as ECM organization, integrin cell-surface interactions, interleukin and interferon signaling, or those associated with the cell cycle, compared to the human TB granulomas. These results collectively indicated that our system significantly recapitulated the Mtb-mediated *in vivo* pathway regulations.

**Fig. 3.**
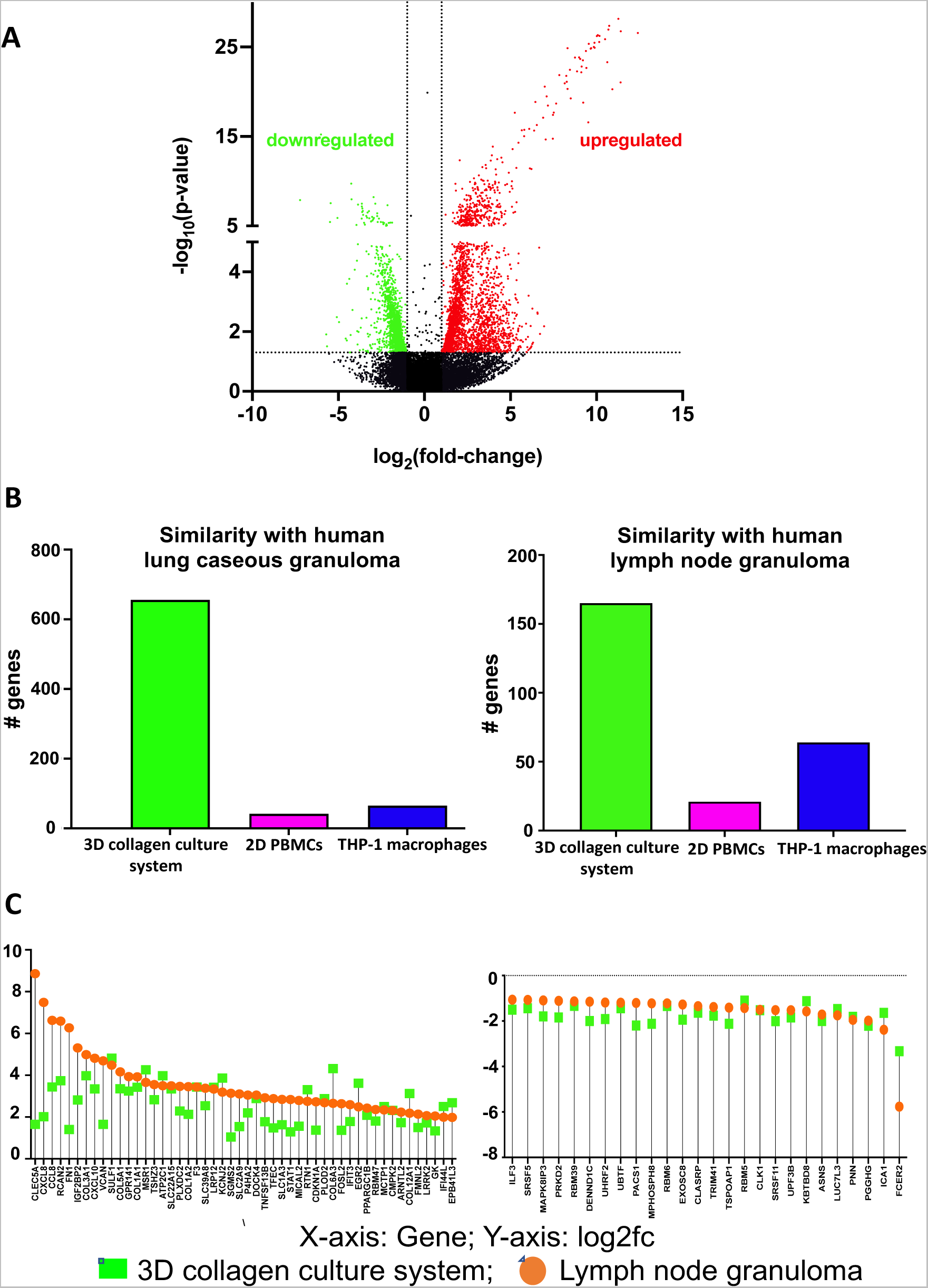
RNA-sequencing of the mammalian cells from the infected collagen gels show maximum similarity with *in vivo* systems. (**A**) A volcano plot of differentially expressed THP-1 genes between uninfected hydrogels and the Mtb infected hydrogels (red: significantly upregulated, green: significantly downregulated, black: non-regulated) (**B**) Quantification of common genes between the various models and the primary TB granulomas (green: 3D collagen culture system, pink: 2D PBMCs, blue: 2D THP-1 macrophages). (**C**) Extent of regulation of the top upregulated genes and downregulated genes common to lymph node granuloma and the 3D collagen culture system.

The transcriptomic analysis of Mtb from the infected collagen gels identified 460 significantly differentially regulated genes (|log2fc|>1, FDR<0.05, 241 upregulated and 219 downregulated, **Supplementary Table 4**) when compared to the planktonic bacteria cultured in 7H9 medium. Functional categorization revealed the upregulated genes involved in crucial infection-related functions such as virulence and adaptation and cell wall processes, among others (**Supplementary Fig. S7**). Given the lack of robust Mtb transcriptomic data from human infections, the comparisons were made with observations from infected mice (9), which showed 114 genes to be similarly regulated in our hydrogels (55 genes upregulated and 59 genes downregulated) (**Fig. 4A**, **Supplementary Table 5**). Mtb transcriptome from THP-1 macrophages infected for 24 h in 2D (54) showed only 21 genes (all upregulated) to be similarly regulated compared to mice infection (**Fig. 4A**, **Supplementary Table 5**). This Mtb transcriptomic signature in 2D (from early time points as compared to 3D) pointed towards the pathogen’s adaptation to the mammalian cells, wherein, genes associated with zinc ion homeostasis, type VII secretion system, and membrane repair were significantly upregulated. Our analyses also revealed a large variation between the Mtb transcriptomes from 2D infected cells (54) and the collagen hydrogels, which could probably be attributed to the time spent by the pathogen in the host (24 h in 2D vs 14 days in collagen gels, **Supplementary Table 5**).

**Fig. 4.**
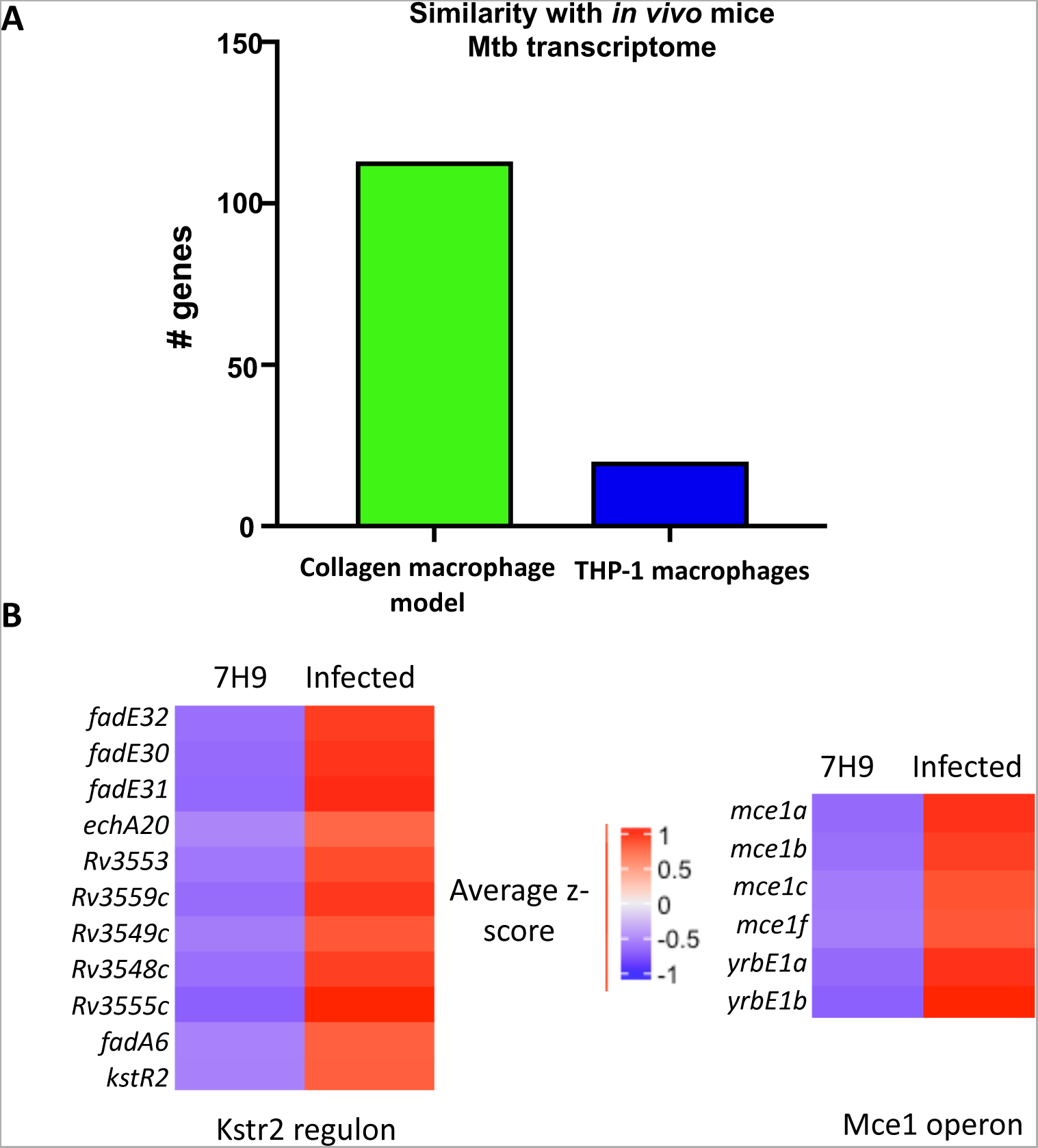
RNA sequencing analysis of bacteria from collagen culture systems shows similarity with mice Mtb transcriptome. (**A**) Quantification of common genes between the *in vitro* models and *in vivo* Mtb transcriptome (GSE132354) in mice (green: 3D collagen culture system, blue: 2D THP-1 macrophages – GSE6209). (**B**) Heat maps for the normalized z-scores of the genes involved in Kstr2 regulon and the mce1 operon. source: Mycobrowser, École Polytechnique fédérale de Lausanne). Genes were considered differentially expressed based on |log2fc| ≥1 and FDR of ≤ 0.05.

We then looked at the functional analyses of the Mtb transcriptome in the model and found significant upregulation of most of the genes of the Kstr2 regulon (**Fig. 4B**), indicating high cholesterol-associated metabolism of the pathogen (9). Intriguingly, we found the upregulation of the mce1 operon genes (*mec1A, mce1B, mce1C, mce1F, mce1R*) (**Fig. 4B**), which are similarly regulated *in vivo,* but oppositely regulated in traditional 2D infection cultures. Kstr2 regulon genes are involved in modulating host response to facilitate persistence (55) and fatty acid transport (56, 57). Moreover, we also found upregulation of genes involved in lipid degradation (*lipL, plcC*), TCA cycle, and fatty acid β-oxidation (*icd1, Rv0893c, echA20*), highlighting the dependence of Mtb on cholesterol and fatty acid utilization in our model, a characteristic of the pathogen’s metabolism in the interstitial and alveolar macrophages, respectively, *in vivo* (9, 35). Furthermore, we also found upregulation of genes of the tryptophan synthase operon (*trpA, trpB, trpC*). Mtb has the ability to synthesize tryptophan, thus counteracting the attempts by T-cells to starve the pathogen of this essential amino acid (58). These results collectively indicate a robust gene signature similarity of Mtb from our system with that from established *in vivo* infection models, while highlighting the drawbacks of using conventional 2D systems for functional studies. Overall, the 3D collagen culture system was able to capture multiple mechanisms underlying Mtb infection-associated host-pathogen dynamics significantly better than the current *in vitro* platforms.

### 4. THP-1 cells accumulate lipid bodies in response to infection

Next, we focused on deciphering the Mtb-mediated metabolic alterations. Mtb can persist in the host for long durations and can switch the energy source required for its metabolism (59) (60). In the initial stages of infection, the pathogen relies on glucose and fatty acids for its metabolism; however, as the infection progresses, fatty acids and cholesterol become the primary energy source (59). Thus, Mtb induces metabolic reprogramming in the macrophages to accumulate triglycerides, leading to the formation of foamy macrophages, a hallmark of human primary granuloma (60). In a 2D system, owing to the shorter duration of infection, the increase in the accumulation of lipids by the infected mammalian cells is not significant **(Supplementary Fig. S8)**. To see if similar to *in vivo* infection conditions, foamy macrophages were generated in our 3D culture system, the uninfected and infected collagen gels were stained for neutral lipids with BODIPY (61). The cells in the infected gels showed a higher abundance of lipid droplets than uninfected gels (**Fig. 5A-B**). Infected gels had a 3-4 folds higher signal on days 7 and 10 post-infection compared to uninfected gels (**Fig. 5C**). Additional confirmation by flow cytometry corroborated these results (**Fig. 5D**). Upon further quantification of BODIPY signal in the infected gels we observed a higher intensity of the signal in the infected cells w.r.t the bystander cells (**Fig. 5E**). Interestingly the BODIPY signal in the bystander cells in the infected gels was still significantly higher than cells from uninfected gels, thus indicating that the bystanders were also getting primed with lipid bodies, indicating a perturbation of microenvironment in the gels due to Mtb infection (**Supplementary Fig. S9**). Transcriptomic data from human caseous granuloma (GSE20050) showed upregulation of 55 genes involved in lipid metabolism (21), and nearly a third of these genes were also upregulated in our culture system; this is in contrast to the negligible overlap with the current *in vitro* 2D models (GSE162729 – THP1 macrophages infected with Mtb in 2D – 1 gene in common; GSE174566- PBMCs infected with Mtb – none), further validating the efficiency with which our system mimics *in vivo* infections (**Fig. 5F**).

**Fig. 5.**
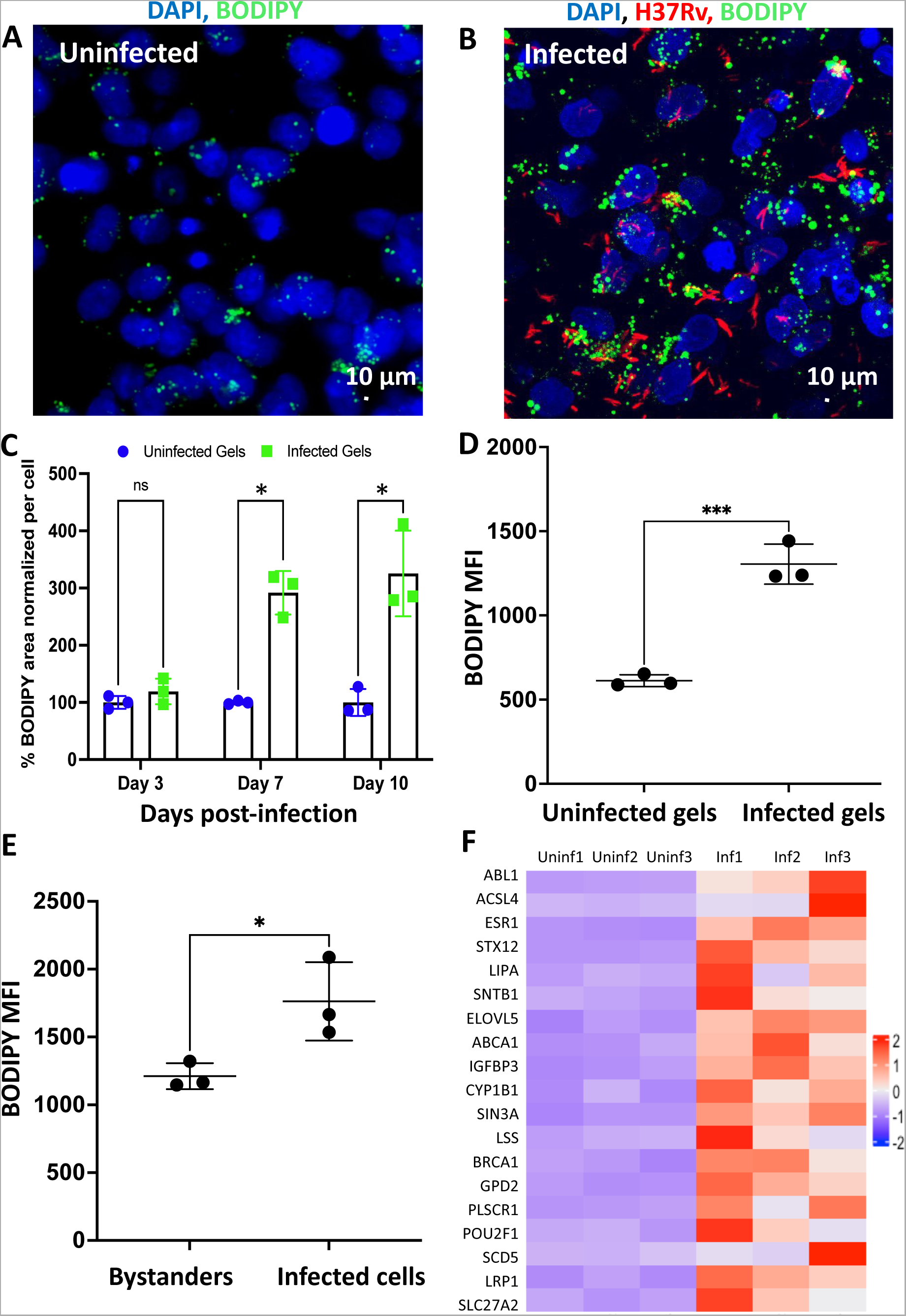
Cells in the infected collagen gels show accumulation of lipid bodies and differentiate to macrophages. Fluorescence microscopy images of uninfected gels (**A**) and gels infected with tdTomato expressing Mtb (red) (**B**) that were stained with the nuclear stain DAPI (blue) and BODIPY (green), 10 days post-infection. (**C**) Quantification of BODIPY staining using ImageJ showed infected gels with a higher stain area per mammalian cell (*n* = 3). (**D**) Mean fluorescence intensity (MFI) of BODIPY signal of THP-1 cells from uninfected gels (Day 10) and infected gels (Day 10) quantified using flow cytometry (**E**) Mean fluorescence intensity (MFI) of BODIPY signal of bystanders and Mtb infected THP-1 cells from infected gels quantified using flow cytometry. (**F**) Heat map of the average z-scores, showing the list of genes involved in lipid metabolism (common with a human caseous granuloma). Data in graphs represent the mean ± s.d., and *p* values were determined by two-tailed unpaired t-tests with Welch’s correction for **C** and without Welch’s correction for **D** & **E** using GraphPad Prism software. \**p*-value < 0.05 was considered significant. Day 7 (\**p*=0.012) and Day 10 (\**p*=0.026) for **C**, *** *p*=0.0006 for **D**, * *p*=0.0348 for **E**

### 5. Pyrazinamide efficacy at or below clinically relevant concentration

Pyrazinamide (PZA) has been a mainstay of antituberculosis treatments since the 1950s and played a major role in shortening the therapy from 12 months to 6 months (62, 63). However, remarkably, it shows no significant activity in any *in vitro* model at clinically relevant concentrations (64, 65), and its anti-tubercular activity was discovered directly on an animal model of infection (66, 67), highlighting how the current *in vitro* culture systems are not optimal for drug testing. Given the exclusive effectivity of pyrazinamide *in vivo*, it has been difficult to fully understand its mechanism of action, and it is only in the recent past that some insights have emerged (14, 68-72).

Our study found that PZA consistently reduced the bacterial burden in our culture system at 50 µg/mL (**Fig. 6A & 6B)**, which is in the range of average peak plasma concentration (C_max_) in individuals undergoing TB treatment with PZA (73). We further validated this finding using primary human monocytes isolated from human PBMCs (**Fig. 6C**). However, at this concentration, PZA failed to act in 2D infection models (**Supplementary Fig. 10A**) and was also ineffective on planktonic bacteria and bacteria embedded in gels without mammalian cells (**Supplementary Fig. S10B-C**). A previous study using collagen-alginate matrix had shown PZA efficacy, but the concentration of the drug used was 500 µg/mL, ten times higher than C_max_ (74). Furthermore, we carried out a dose-response-based study to check the concentration at which PZA becomes effective and observed a significant CFU reduction even at 10 µg/mL of the drug concentration (**Fig. 6D**). The similar efficacy of pyrazinamide between our 3D collagen model and in *in vivo* conditions indicated the potential of our system as a more reliable drug testing platform.

**Fig. 6.**
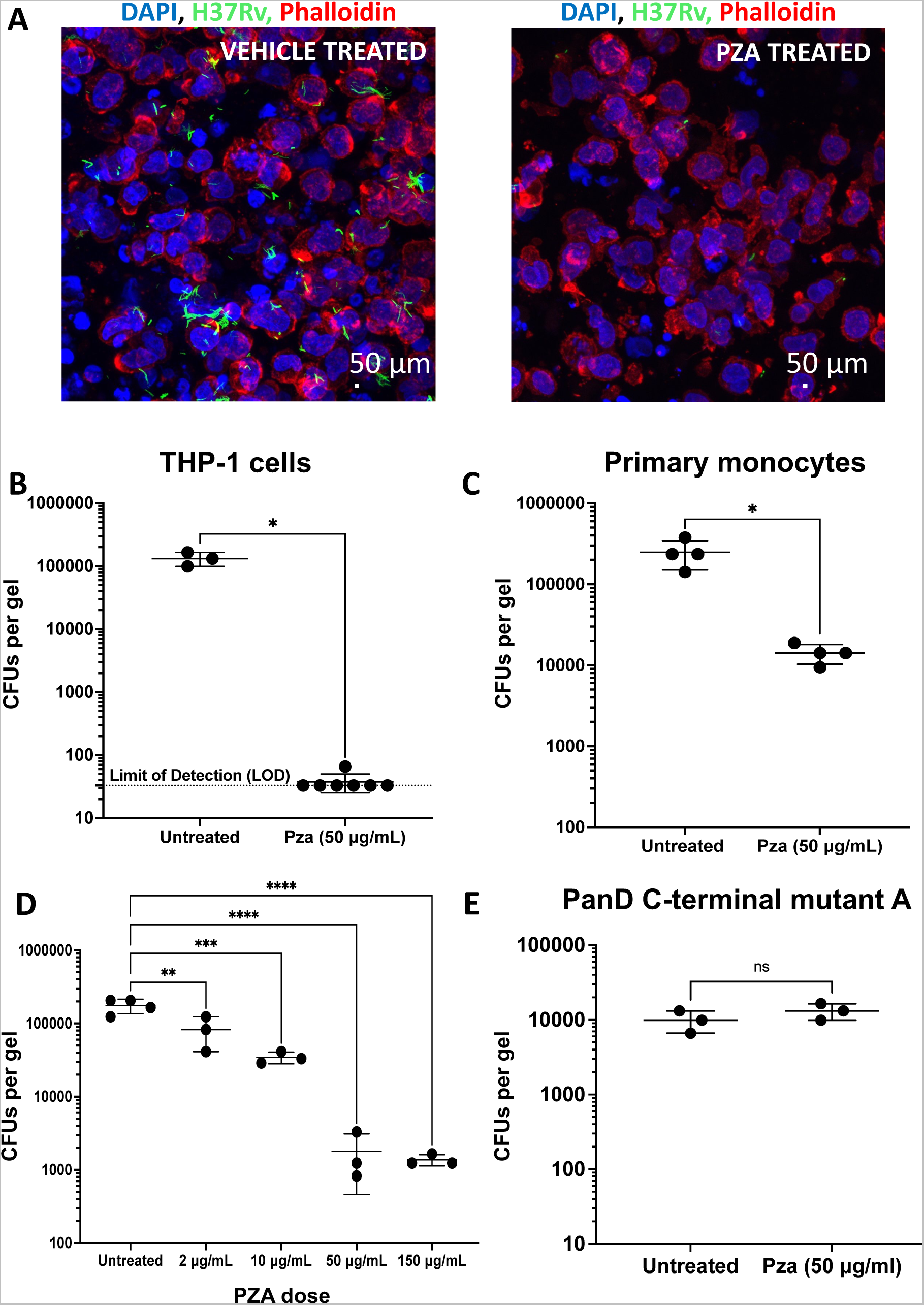
Pyrazinamide showed efficacy at a clinically relevant concentration in infected collagen gels. (**A)** Representative images of sections from Mtb infected THP-1 in collagen gels treated with PZA at 50 μg/mL (right) with respect to the vehicle-treated control (left). In these gels, the infection was carried out for 4 days, post which the experimental groups were treated with PZA at 50 μg/mL for 6 days. Intracellular CFU plots obtained after treating (**B**) Mtb-infected THP-1 in collagen gels (*n* = 3 for untreated and *n* = 7 for treated), the treatment with PZA was started 4-days post-infection and was carried out for 6 days; hence the CFUs shown are 10 days post-infection (**C**) Mtb infected primary human monocytes (*n* = 4), the treatment with PZA was started 7-days post-infection and was carried out for 3 days; hence the CFUs shown are 10 days post-infection. **(D)** Intracellular CFU plots were obtained after treating Mtb infected THP-1 in collagen gels (*n* = 4 for untreated and *n* = 3 for other groups), the treatment with PZA was started 4-days post-infection and was carried out for 6 days; hence the CFUs shown are 10 days post-infection. **(E)** Mtb (POA resistant strain) infected THP-1 cells with PZA at 50 µg/mL (ns: non-significant) (*n* = 3), the treatment with PZA was started 4-days post-infection and was carried out for 6 days; hence the CFUs shown are 10 days post-infection. The mutations are listed in Supplementary Table 6. Data in graphs represent the mean ± s.d., and *p* values were determined by two-tailed unpaired t-tests with Welch’s correction for **B&C** and Tukey’s multiple comparison tests for **D** using GraphPad Prism Software. *p*-value < 0.05 was considered significant. \**p*=0.0202 for **B** and \**p*=0.0171 for **C;** \**p*=0.0068 for **untreated vs 2 μg/mL,** \**p*=0.0002 for **untreated vs 10 μg/mL,** \**p*<0.0001 for **untreated vs 50 μg/mL,** \**p*<0.0001 for **untreated vs 150 μg/mL** for **D,** ns = non-significant for **E**

Pyrazinamide is a prodrug catalyzed to its active form pyrazinoic acid (POA), by the bacterial gene *pncA* (75). One of the working hypotheses of PZA points to its ability to perturb the intrabacterial pH (76). Briefly, PZA gets metabolized to POA inside the bacteria and is transported out through the efflux pumps. However, in an acidic pH microenvironment (such as in a phagosome), POA gets protonated to HPOA, which then re-enters the bacteria. This cycle eventually leads to the bacteria being unable to maintain its cytoplasmic pH and membrane potential, leading to death. We tested this hypothesis by checking for phagosomal maturation in the infected 2D and 3D conditions and did not find any significant differences in colocalization of LAMP-1-stained lysosomes with Mtb in the 2D infection scenario compared to 3D (**Supplementary Fig. S11**). We also compared the Mtb genes from the 3D collagen culture system, THP-1 macrophages (2D infection - GSE6209), to those induced by low pH as an *in vitro* stimulus (77). This analysis showed 16 genes of Mtb in the 3D collagen culture system to be common with those induced by low pH as against 19 genes of Mtb from the 2D infection, further pointing to relatively lower/similar pH stress in the 3D collagen culture system as against 2D infection (**Supplementary Table 6**). These results collectively indicated that the pH-dependent effectivity of PZA is not the major mechanism in our model.

PZA resistance in clinical samples is not always associated with mutations in *pncA* (78–80). A few studies have pointed to the critical role of Aspartate Decarboxylase *panD* and the Unfoldase *clpC1* genes in mediating resistance to PZA (70, 71). PanD is involved in the synthesis of pantothenic acid (vitamin B5), critical for biosynthesis of coenzyme A and acyl carrier protein (ACP), involved in various steps of lipid metabolism, and hence bacterial mutants with deletion of these genes were found to be highly attenuated (81). The active form of the drug, POA, binds to PanD and the C-terminal end of the PanD-POA complex is recognized by the caseinolytic protease, ClpC1-ClpP, leading to its degradation (68). We generated four spontaneous POA mutants of H37Rv (labelled mutant A to D) as previously described (70) and corroborated that mutations were present in the *panD* gene corresponding to the C-terminal domain of PanD (**Supplementary Table 7**). *pncA* and *clpC1* genes did not have any mutations in these bacteria. We infected THP-1 cells with these POA mutants in the collagen gels and treated the experimental groups with PZA (50 µg/mL) (**Fig. 6E**, **Supplementary Fig. S12A-C**). PZA was ineffective in reducing the bacterial burden for all 4 mutants, clearly emphasizing the critical role of PanD in the efficacy of PZA in the 3D model and strengthening the case for adopting this culture system in drug discovery and mechanistic studies.

Several other proteins are hypothesized to synergize/antagonize PZA activity (82, 83). We found that these Mtb genes were not differentially regulated in our gels compared to planktonic cultures (**Supplementary Table 8**). Thus, the transcriptomic data suggest that these proteins may have a minor role in PZA efficacy in our culture system, although there may be differences at the translational level. Further validation of the system was done using the other first-line anti-TB drugs (rifampicin, ethambutol, or isoniazid), which were also effective in reducing bacterial counts when used at their respective minimum inhibitory concentrations (MIC) of broth culture (**Supplementary Fig. S13 A-C**).

## Discussion

Tuberculosis causes a significant social and economic burden on both patients and society. There is a need for active research to understand the bacteria-host interaction and find new drug candidates. Although experiments on planktonic cultures of Mtb and 2D infection experiments with mammalian cells continue to be the primary step for many studies, their inherent limitations often lead to findings that do not translate well to animal models and humans. Such efforts consume precious time and resources or could also curtail the development of effective therapies, warranting the need to develop better drug-testing platforms. In this study, we propose a 3D collagen-hydrogel culture system as a next-generation *in vitro* platform to delineate the complexities of Mtb infection by mimicking an *in vivo* system. 3D culture platforms are being extensively used to study many diseases and are gaining wide acceptance (18). The current culture system uses hydrogels made of collagen 1, the main ECM component of human lungs. ECM plays a fundamental role in TB pathogenesis during human infection, as Mtb is known to induce the production of matrix metalloproteases (MMPs), which digest collagen, leading to ECM degradation and disease progression (84). Hence, inhibitors that act on these proteases are being studied as potential supplements to the existing treatment regimens (85). The 2D infection studies lack ECM, and mice are not known to produce orthologs of some MMPs found in humans (86). Among the various 3D platforms, a few studies have cultured peripheral blood mononuclear cells (PBMCs) with Mtb in a three-dimensional (3D) matrix to form micro aggregate structures of cells of nearly 100 µm surrounding an infected cell or bacteria (87) and showed that collagen reduced mammalian cell death upon Mtb infection and Mtb growth was arrested in such systems (20, 88). Primary granulomas from zebrafish have also been cultured *ex vivo* (89). More advanced 3D systems include engineering lung organoids using stem cells, though its potential is currently unexplored in TB research (10). A recent study has used human airway organoids to study mycobacteria interaction with airway epithelial cells (90). Furthermore, *in vitro* hollow fibre system for TB (HFS-TB) was used to recapitulate the human pharmacokinetics and pharmacodynamics of a few TB drugs, which resulted in optimal dosing regimens for moxifloxacin and linezolid (91). Although all these platforms add significant value to the field, there are difficulties in procuring primary lung granulomas, lung biopsies, and specialized equipment in BSL-3 containment. Furthermore, none of these systems has shown the efficacy of pyrazinamide at clinically relevant concentrations.

Hence, given the presence of natural ECM components like collagen, a suitable mechanical environment, and 3D culture, we believe the 3D system proposed here is amenable to widespread adoption and will significantly contribute to testing and validating new hypotheses for transitioning from *in vitro* to *in vivo*. Collagen gels are easy to synthesize, widely used for many applications (92), and easy to work with even in BSL-3 facilities. These collagen gels have been extensively characterized and are modular, for example, the matrix characteristics, such as pore size, stiffness, etc., depend on polymerization temperature and can be tuned in accordance with the human lung ECM (93).

THP-1 monocytes and BMDMs are commonly used for studying host-pathogen interaction *in vitro*, but the length scale of these studies continues to be a major challenge. BMDM infection offers a slightly longer duration to study host-pathogen interaction, and a previous study had characterized the transcriptional and physiological profile of Mtb in infection with BMDMs for 14 days showing the adaptation of the pathogen in the face of host stress (38). However, the kinetics of the BMDMs response in terms of viability varies greatly with MOI, with approximately 75% at day 6, with an MOI of 1, which falls to less than 25% with an MOI of 5 (94). Our culture system also offers the flexibility of looking at host-pathogen interaction at multiple levels such as time (1 day – 3 weeks) and scale: single cell to multiple cells behaviour for mammalian and Mtb; and is suitable for analysis through tools such as genetic mutations in bacteria and host, histology, fluorescence microscopy, CFU assays, live-cell imaging, and flow cytometry.

We also show a genotypic and phenotypic similarity of mammalian cells in the infected collagen gels to human infection. For example, Mtb is known to induce the formation of lipid bodies in macrophages *in vivo* (21, 95). Previously, it has been shown that 2D infection of human macrophages can lead to a high accumulation of lipid bodies compared to uninfected macrophages (96), however, more recent reports indicate necrosis of other cells in the milieu to be critical for this phenomenon. An MOI of 1 or 5 in 2D cultures did not result in foamy macrophage formation but instead was seen with a high MOI of 50, which led to more than 60 % mammalian cell death (97). The necrotic cells served as a source of triglycerides for the live infected cells. Other commonly used methods include supplementation of oleate in the growth media to generate foamy macrophages, followed by infecting these cells with Mtb to study lipid retention/turnover (98, 99). Another model used PBMCs exposed to sepharose beads coated with Mtb purified protein derivative (PPD) to show aggregate formation accompanied by foamy macrophages (100). Mtb infection of murine adipocytes (101) or supplementation with lipoproteins (102) have also been tried to induce the formation of foamy macrophages. Furthermore, human macrophages differentiated in 2D have a propensity to accumulate lipids even in the absence of Mtb infection (21). This could explain the absence of an increase in foamy macrophages after infection in our 2D infection, as we used a low MOI of 1, which is unlikely to induce significant necrosis. In our collagen culture system, even a low burden of Mtb infection led to this phenotype without any additional supplementation of fatty acids. We also show cord-like Mtb replication in the infected collagen gels, another prominent phenotype found *in vivo* (22, 23, 47, 48). Furthermore, in our Mtb transcriptomic data, upregulation of the gene *fbpC*, a member of the antigen 85 family of proteins that confer fibronectin binding affinity and maintain the cell wall integrity of Mtb by synthesizing alpha trehalose-dimycolate (TDM, cord factor) (103), elucidated the molecular basis of cording in our system. Host cell autophagy is an important defense mechanism against intracellular pathogens, and the upregulation of autophagic genes and pathways in our dual RNA sequencing data points towards such a phenomenon also occurring in our system, which the bacteria successfully evade by cording (50). Transcriptomic analysis of the mammalian cells in the infected collagen gels revealed a significantly greater overlap with human TB granulomas than in current culture systems. Such shortcomings in how mammalian cells in 2D respond to Mtb infection render them somewhat unreliable systems for testing potential drug candidates. For example, none of the current *in vitro* systems could demonstrate significant PZA efficacy at clinically relevant concentrations (≤ 50 µg/mL), which could be observed using our 3D collagen culture system. PZA has been shown efficacious *in vitro* using planktonic bacteria at a pH of 5.95, with a MIC of 200 µg/mL (76). Further reduction in pH of medium resulted in growth inhibition of several strains of bacteria (76). Similarly, in human monocyte-derived macrophages, treatment with concentrations as high as 100-400 µg/mL of PZA only resulted in about 70% reduction in bacteria as determined by fluorescence imaging (72, 104). Thus, our platform is potentially an excellent substitute for current *in vitro* models and a starting point for more accurate drug testing and to give deeper insights into host-pathogen interaction before large-scale animal studies.

The tuberculosis bacilli in a host are taken up by monocytes, macrophages, dendritic cells, and neutrophils (7) and also have a niche in mesenchymal stem cells and lymphatic endothelial cells (50, 105). Also, since we wanted to mimic natural infection, PMA was not used to differentiate THP-1 cells. In our culture, PZA was also efficacious when used on Mtb-infected primary human monocytes. However, other observations regarding mammalian and bacterial phenotypes need to be validated with primary cells. Another limitation is that the culture system does not incorporate other immune and non-immune cells that are relevant *in vivo.* During infection, Mtb primarily resides in alveolar macrophages (AM) and interstitial macrophages (IM) *in vivo* and has distinct interactions with each of these macrophage populations (9); Ams offer a more permissive niche for bacterial growth compared to IMs (35), and hence, it would be interesting to see how AMs respond to infection in our collagen hydrogels. However, there are no established human AM cell lines, and primary AMs are difficult to obtain. Furthermore, the bacteria encounter multiple stresses *in vivo,* such as adaptive immune response and hypoxia in a necrotic granulomatous core, which was not present in our platform. The absence of other components of the adaptive immune system also means that Mtb can continue to proliferate in the macrophages as the duration of infection progresses. It is possible to increase the density of cells cultured in collagen gels, which may lead to low oxygen concentration near the center of the gel, as shown in spheroid cultures (106) and will be explored in future studies.

## Conclusions

We show that our 3D collagen cell culture model offers a significantly longer window for studying host-pathogen interaction. Cells in the infected microenvironment have properties similar to cells in *in vivo*, in terms of lipid accumulation, cord formation, and gene expression, among other characteristics. The effectivity of pyrazinamide, a first in any *in vitro* model, in reducing the bacterial burden further emphasizes the robustness of the platform, and we hypothesize that potentially undiscovered mechanistic phenomena can be unravelled using our culture system. The system offers utility with ease of usage and modularity while incorporating resources that are easily accessible to the research community and can therefore be readily adopted.

**Supplementary video 1: Z-stack of infected collagen gel (DAPI – Blue, H37Rv – Green, Phalloidin – Red).** The video shows different planes of the collagen gel over time. (Step size – 0.5 μm, total depth of view – 79 μm)

**Supplementary video 2: 3D view of infected collagen gel (DAPI – Blue, H37Rv – Green, Phalloidin – Red)**

## Methods

### 1. Culture of mammalian cells and isolation of primary human monocytes

Human THP-1 monocytes were seeded at a concentration of 10^5^ cells/mL and passaged every 3 days. The cells were cultured in RPMI 1640 (Invitrogen) media containing 10% Fetal bovine serum (FBS) and antibiotics (Penicillin/Streptomycin).

For human primary monocytes, 10 mL of whole blood was collected from volunteers (Institutional Human Ethics Committee Approval Number: 20-14012020) and peripheral blood mononuclear cells (PBMCs) were isolated by density gradient centrifugation over Histopaque^®^-1077 (Sigma Aldrich). From these PBMCs, the primary monocytes were isolated using a monocyte isolation kit (Mojosort^TM^ Human CD14+ Monocytes Isolation Kit, BioLegend^®^), per the manufacturer’s instructions. These primary monocytes were cultured in RPMI 1640 media supplemented with 10% human serum (H4522, Sigma Aldrich).

All mammalian cells were cultured in a humidified incubator supplemented with CO_2_ and maintained at 37 ^°^C.

### 2. Culture of H37Rv

*Mycobacterium tuberculosis* strain H37Rv was cultured in media composed of 7H9, tween 80 (0.05%), glycerol (0.1%) with albumin-dextrose-catalase (ADC) (10%) enrichment at 37 °C. GFP-H37Rv carrying the plasmid pmyc1tetO-gfp, tetR(B), and tdTomato H37Rv carrying the plasmid pTEC27 (pMSP12: tdTomato), with hygromycin (50 μg/mL) as the selection marker, were used for imaging and flow cytometry experiments. Mtb H37Rv-pBP10 was cultured similar to above with 30 μg/mL kanamycin.

### 3. Collagen gel preparation

Rat tail Collagen I (3 mg/mL, Gibco^TM^) was used. Collagen gels with final concentrations of 1 mg/mL to 2 mg/mL were made as per the manufacturer’s recommendations. The pH of these gels ranged from 7-7.4. The polymerization was carried out in a humidified incubator at 37 °C for 20 minutes, following which they were transferred into a 24-well plate containing 1X phosphate-buffered saline (PBS). In experiments involving mammalian cells and bacteria, the cell suspension (4×10^6^ cells/mL) was added in place of 1X PBS, and the gels were transferred to wells containing RPMI 1640 containing 10% FBS. Each collagen gel had 2×10^5^ cells and was 50 μL in volume with a diameter of approximately 2.5 mm.

### 4. Infection experiments

Infected collagen gels were made as follows: single-cell suspension of H37Rv (OD of 0.5-1) was prepared and added to THP-1 monocytes at a multiplicity of infection (MOI) of 40:1 (bacteria: mammalian cells) in the required volume for making the gels. Post gelation, the infection was carried out for 12 h, after which amikacin sulfate (200 µg/mL) was added to the media for 6 h to remove extracellular bacteria. The media was changed every 48 h during the experiment. The gels were treated with collagenase, type 1 (0.4 mg/mL, GIBCO™) for 1 h at 37 °C to degrade the collagen for flow cytometry experiments. The cells were stained with LIVE/DEAD^TM^ (L34955 LIVE/DEAD™ Fixable Violet Dead Cell Stain Kit, for 405 nm excitation), followed by fixation with 4% paraformaldehyde (PFA), and used for further analysis.

For 2D infection experiments, THP-1 cells were differentiated into adherent macrophages using 20 nM Phorbol 12-Myristate 13-Acetate (PMA). Briefly, cells were incubated in media containing PMA for 24 h, post which the PMA was removed, and the cells were allowed to rest for 2 days with fresh media. Cells were then infected with Mtb at an MOI of 1:1 (bacteria: mammalian cells) for 4 h and treated with amikacin sulfate (200 μg/mL) for 2 h.

For infection in collagen gels, the primary human-derived monocytes were used without further differentiation, and the infection at an MOI of 1 was carried similarly to THP-1 cells as described above.

### 5. Mtb growth kinetics

To study the kinetics of Mtb replication during infection in the hydrogel system, we infected THP-1 cells with H37Rv carrying the replication clock plasmid, pBP10 (39) (a kind gift from Dr. David Sherman). Intracellular CFUs were carried out by treating the gels with collagenase, lysing the mammalian cells with 0.1% Triton X-100, followed by spotting serial dilutions on Middlebrook 7H11 agar plates supplemented with 10% OADC, with and without 30 µg/mL kanamycin and incubated at 37 °C for 20-24 days. The replication and death rates of Mtb were quantified using the mathematical model proposed by Gill et al.(39).

### 6. Second-harmonic imaging of collagen gels and lung tissues from mice

Second-harmonic generation (SHG) images of the samples were acquired using a mode-locked fiber laser (Coherent Fidelity HP) with an operational wavelength of 1040 nm, a pulse width of 140 fs, and a repetition rate of 80 MHz. The incident beam was scanned using a galvo-scanner (GVS001, Thorlabs), with the beam focused on the sample using a 60x/1.2 NA water immersion objective (UPLSAPO60XW, Olympus). A photomultiplier tube (R3896, Hamamatsu) was used to collect the SHG emission at 520 nm from collagen. The incident power at the focus for imaging collagen was 1.7 mW. The average pore size of the collagen gels was calculated using the particle analyzer in the BoneJ plugin of ImageJ (NIH, USA). The resolution of the microscope was around 400 nm, and hence the pores below the size of 0.16 µm^2^ were excluded from the analysis.

### 7. Atomic Force Microscopy (AFM) measurement

Collagen gels were loaded into custom polydimethylsiloxane (PDMS)-based AFM holder. The sample was kept hydrated during measurement, and indentation (with a depth of 500 microns) was done at different points using Park Systems NX-10 AFM in a liquid medium. Force distance spectroscopy was carried out using the HYDRA6V-200NG-TL cantilever with a 5 µm diameter sphere. The force constant was 0.045 N/m, and the force limit was 0.91 nN. The up and down speed was 0.3 µm/s. The data generated were further analyzed through a custom-written MATLAB code to determine Young’s modulus (30).

### 8. Ferret diameter calculation of Mtb cords

THP-1 cells were infected with GFP-H37Rv in collagen gels for 10-14 days, post which the gels were fixed with 4% paraformaldehyde (PFA). The gels were stained with Hoechst 3342 (Thermo Scientific™) and Phalloidin (A12380, Thermo Scientific™) and imaged using a confocal microscope (SP8, Lecia Microsystems). The GFP signal from H37Rv from 5 images was thresholded, and only aggregates above 5 μm^2^ were used for further analysis to filter out the single bacteria/noise. The ferret diameter was then calculated using the Analyze option in ImageJ.

### 9. TB antibiotic treatments

THP-1 cells were infected with Mtb in collagen gels for 7 days, post which the gels in the treatment group were treated with the first line of antituberculosis drugs – rifampicin (Himedia^®^) (0.3 μg/mL), isoniazid (Himedia^®^) (0.25 μg/mL), and ethambutol (Himedia^®^) (5 μg/mL). At 3- and 7-days post-treatment, gels were degraded with collagenase, and cells were lysed with 0.1% Triton X-100. Cell lysates were plated on 7H11 plates supplemented with 10% OADC and incubated at 37° C. The CFUs were counted 3-4 weeks after plating. For studying pyrazinamide (Himedia^®^) (50 μg /mL) efficacy, the infection was carried out for either 4 or 7 days, post which the cells were treated for 6 or 3 days, respectively.

### 10. Generation of pyrazinoic acid (POA) mutants

The H37Rv POA mutants were generated as described by Pooja et.al. (68). Briefly, Mtb was grown on 7H11 plates supplemented with 2 mM POA and 10% OADC, and a few colonies were picked and restreaked on fresh 2 mM POA plates. Colonies from 2^nd^ plate were further grown in 7H9 media supplemented with 2 mM POA and 10% ADC. Genomic DNA was isolated from these mutants and sequenced with gene-specific primer pairs (**Supplementary Table 9**) for *pncA, panD, clpC1* genes using Phusion^TM^ High-Fidelity DNA polymerase (Thermo Scientific^TM^) as per the manufacturer’s instructions. The products were sequenced using Sanger sequencing and the results were analyzed using the NCBI BLAST tool.

### 11. Antibody & BODIPY staining

Differentiation of monocytes to macrophages was quantified by CD68 expression. Briefly, cells in the infected and uninfected hydrogels were released using collagenase (0.4 mg/mL). The cells were stained with LIVE/DEAD^TM^ (L34955 LIVE/DEAD™ Fixable Violet Dead Cell Stain Kit, for 405 nm excitation), post which they were fixed using 4% PFA. The cells were then permeabilized using 0.1% Triton X-100 and stained with anti-CD68 antibody (PE-Mouse Anti-Human CD68 set, BD Biosciences) (1:50 dilution) overnight in a buffer made of 1X PBS supplemented with 1% bovine serum albumin (BSA).

To study the activation of the THP-1 cells post-infection, cells were stained with an anti-iNOS antibody (NB300 – Novus Biologics). Briefly, at day 10, cells in the infected and uninfected hydrogels were released using collagenase (0.4 mg/mL). The cells were stained with LIVE/DEAD^TM^ (L34955 LIVE/DEAD™ Fixable Violet Dead Cell Stain Kit, for 405 nm excitation), post which they were fixed using 4% PFA. The cells were incubated in a blocking buffer made of 1X PBS supplemented with 1 % BSA, 0.1% Triton X-100 and 5% donkey serum for two hours at room temperature. This was followed by washing with 1X PBS and staining with the primary antibody (iNOS - 7 μg/mL) diluted in FACS buffer and incubation at 4 °C overnight. This was followed by staining with the secondary antibody (Donkey anti-Rabbit IgG (H+L) Highly Cross-Adsorbed Secondary Antibody, Alexa Fluor™ Plus 647) diluted in the blocking buffer (0.5 μg/mL) for 3 hours, followed by running the samples through a flow cytometer.

To study the localization of Mtb phagosomes with lysosomes in the infected 2D and 3D scenarios, cells were stained with anti-(LAMP-1) antibody (ab24170, Abcam). Briefly, the cells in 2D were incubated in a blocking buffer made of 1X PBS supplemented with 1 % BSA, 0.1% Triton X-100 and 5% donkey serum for two hours at room temperature. The cells were then washed with PBS and stained with the primary antibody (0.5 μg/mL) diluted in the blocking buffer and incubated at 4 °C overnight. This was followed by staining with the secondary antibody (Donkey anti-Rabbit IgG (H+L) Highly Cross-Adsorbed Secondary Antibody, Alexa Fluor™ Plus 647) diluted in the blocking buffer (1 μg/mL), for 3 hours and imaged at 60X using a fluorescence microscope (IN Cell Analyzer 6000). For LAMP-1 staining in hydrogels, the gels were fixed with 4% PFA, post which cryo-sectioning (107) was carried out. The sections were stained with primary and secondary antibodies similar to 2D cultures. For the 2D infection, analysis was done on days 3 and 5 post-infection, while for 3D, it was done on days 5 and 7 post-infection. The images were acquired with a z-stack of 0.5 slice thickness over 20 µm. The co-localization of Mtb and LAMP-1 in the images was analyzed using the Just Another Co-localization Plug-in (JACoP) in ImageJ. Briefly, the intensities of the fluorescence signals (green – Mtb and red- LAMP1-1) were thresholded, and the Pearson’s coefficient from the plug-in was used to calculate the extent of co-localization.

For staining the mammalian cells in 2D and in the hydrogels with BODIPY^TM^ 493/503 (Thermo Scientific™), the cells were fixed with 4 % PFA for 1 hr, followed by incubation with the dye diluted in 1X PBS (1 µg/mL) for 15 mins at 37 °C. The dye was washed, followed by staining with Hoechst 3342 (Thermo Scientific™) and imaged using a fluorescence microscope (IN Cell Analyzer 6000) with a 40X objective. The intensity of the fluorescence signal (green – BODIPY) was thresholded, and the area was calculated using the Analyze Particles option in ImageJ. The obtained BODIPY area per field was then divided by the number of mammalian cells in the field to arrive at the BODIPY area per cell in the field. Analysis was carried out for 10 fields per gel. For flow cytometric quantification of BODIPY, the gels were degraded with collagenase, type 1 (0.4 mg/mL, GIBCO™) for 1 h at 37 °C. These cells were stained with LIVE/DEAD^TM^ (L34955, Thermo Fisher Scientific), followed by fixation with 4% PFA. The cells were then stained with BODIPY, as described above.

For all the flow cytometry experiments, the cells were suspended in 300 μL of FACS buffer, and data were acquired using BD FACSDiva™ software, followed by analysis using FlowJo (v9). The gating strategy to identify the suitable cell populations for CD68, iNOS and BODIPY staining is shown in **Supplementary Fig. S14,** and the viability assay is in **Supplementary Fig. S15.**

### 12. Preparation of samples for dual RNA-sequencing

The gels were degraded using collagenase (0.4 mg/mL) at day 14. The cells were treated with Guanidinium thiocyanate buffer (GTC buffer) containing beta-mercaptoethanol. The homogenate was centrifuged at 5000 g resulting in pellets of Mtb. The supernatant was collected and processed for THP-1 RNA extraction. Mtb RNA was isolated from the pellet using FastRNA® Pro Blue kit (MP Biomedicals™) per the manufacturer’s recommendations. Briefly, 1 mL of RNApro^™^ solution was added to the bacterial pellet and was transferred to a lysing matrix tube. Bead beating was carried using a FastPrep^®^ instrument (2 cycles of 20 seconds each at a speed setting of 4.5 m/s). The isolated RNA was purified using a NucleoSpin RNA kit (MACHEREY-NAGEL) per the manufacturer’s recommendations and quantified using a NanoDrop™ One/One^C^ Microvolume UV-Vis Spectrophotometer (Thermo Scientific^TM^).

### 13. Bioinformatic pipeline and analysis

The libraries were paired-end sequenced on Illumina HiSeq X Ten sequencer for 150 cycles (Illumina, San Diego, USA) following the manufacturer’s instructions. The data obtained from the sequencing run was de-multiplexed using Bcl2fastq software v2.20, and FastQ files were generated based on the unique dual barcode sequences. The sequencing quality was assessed using FastQC v0.11.8 software. The adapter sequences were trimmed, bases above Q30 were considered, and low-quality bases were filtered off during read pre-processing and used for downstream analysis. The pre-processed high-quality data were aligned to the human reference genome using HISAT with the default parameters to identify the alignment percentage. Reads were classified into aligned (which align to the reference genome) and unaligned reads. The tool featureCount was used to estimate and calculate transcript abundance. Absolute counts for transcripts were identified and used in differential expression calculations. DESeq was used to calculate the differentially expressed transcripts. Transcripts were categorized into up, down, and neutrally regulated based on the log2fold change (log2fc) cutoff of ±1 value, and with a false-discovery rate (FDR) < 0.05 were selected for further analysis (GEO accession code GSE216503).

The following publicly available data sets were used for analysis: GSE162729 – for THP-1 macrophages; GSE20050 – for human caseous granuloma, GSE17443 – for lymph node granulomas and GSE174566 -2D PBMCs. Mtb datasets include GSE6209 – for the transcriptional response of Mtb from THP-1 macrophages (2D infection) and GSE132354 – for the transcriptional response of Mtb from mice. From these data sets, genes with a |log2fc|>1 having a false-discovery rate (FDR)<0.05 were selected for further analysis. The genes in the five datasets were compared, and the similarity of the *in vitro* models (3D collagen culture system, THP-1 macrophages, 2D PBMCs) with the primary granulomas (lymph node and caseous) was determined by the number of similarly regulated genes (upregulated and downregulated). The ReactomePA package in R was used for gene ontology enrichment analysis of the differentially regulated genes (108). From the human caseous and lymph node granulomas datasets, the significantly enriched (p<0.05) similarly regulated (upregulated and downregulated) REACTOME pathways in both the data sets were selected for representation, and the regulation of these pathways was compared with the *in vitro* model’s data sets. Heatmaps were made using the ComplexHeatmap function in R.

### 14. Statistical analysis

Data were presented as mean ± standard deviation and plotted using GraphPad PRISM 9 software. Measurements for each data point were taken from distinct samples. Unpaired two-tailed t-tests with or without Welch’s correction were used to detect statistical differences between groups depending on the number of variables, with *p <*0.05 considered significant.

## Acknowledgements

We thank Dr. David Russell (Cornell University) for providing valuable external feedback on this manuscript. We thank Dr. Deepak Saini (IISc) for providing us with GFP and tdTomato conjugated stains of H37Rv and Dr. Amit Singh (IISc) and Dr. Narendra Dixit (IISc) for their invaluable inputs on this project. We thank the Centre for Infectious Diseases Research (CIDR) at IISc Bangalore for allowing us to carry out the studies in the Biosafety Level-3 facility. We thank Dr. Siddharth Jhunjhunwala (IISc) for access to the flow cytometer. We also acknowledge Dr. Thomas Dick’s (Center for Discovery and Innovation, Hackensack Meridian Health) advice for generating spontaneous mutants of PanD. The fluorescence image acquisition was carried out at the Microbiology and Cell Biology (MCB) imaging facility at the Indian Institute of Science.

## Data Availability

The datasets generated during and/or analyzed during the current study are available from the corresponding author on reasonable request. The RNA sequencing data is available at GSE216503.

## Author contributions

RA and VG wrote the manuscript. RA and VG designed experiments. APS helped with the initial optimization of collagen gels. VG performed experiments and analyzed the results. APS and JKM performed the second-harmonic generation and 2-photon imaging. SJ helped with characterizing the POA mutants. VV helped in sample preparation for imaging and validation of PZA data.

