## Supplementary Information for "Lung-mimicking 3-Dimensional hydrogel culture system recapitulates key tuberculosis phenotypes and demonstrates pyrazinamide efficacy"

### Supplementary data

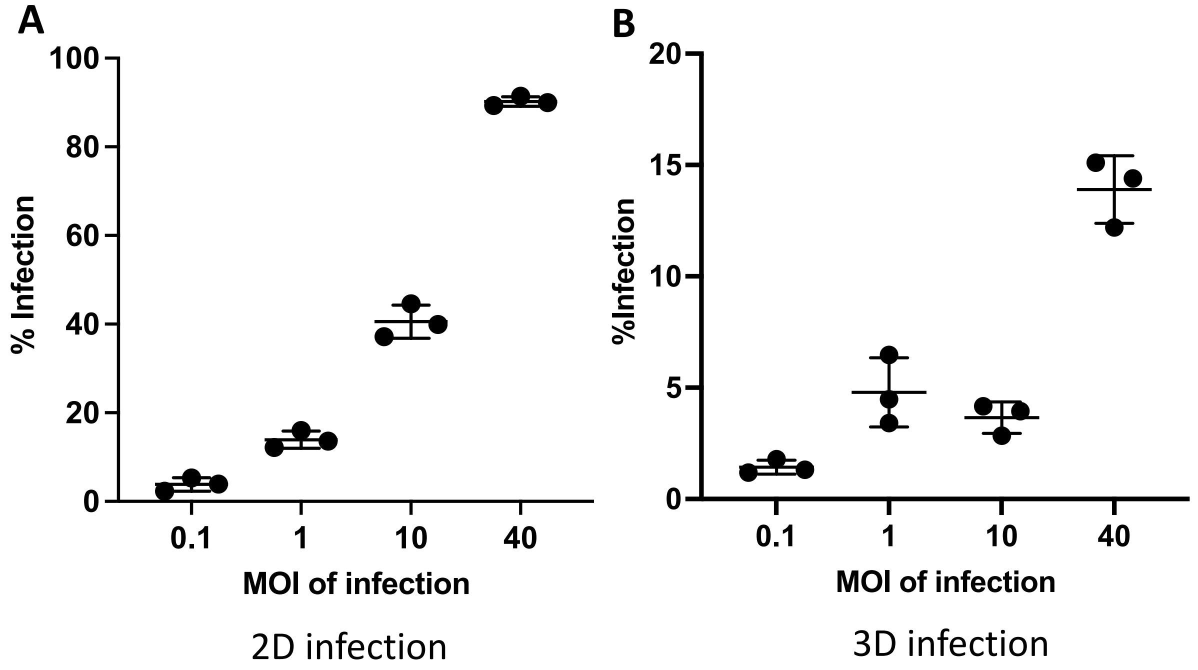


**Fig. S1 Percentage of infected cells can be varied by changing the MOI in both 2D and 3D cultures**. Characterizing the infection % through flow cytometry in 2D (**A**) and 3D (**B**) using a range of MOIs. The infection % with MOI of 1 in 2D infection was in the same range as with MOI 40 in 3D infection (*n* = 3 for each MOI).

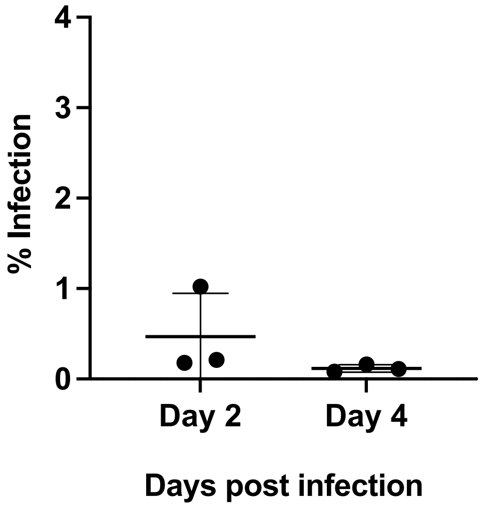


**Fig. S2 Monocyte infection is inefficient without collagen gels.** Infecting THP-1 monocytes with Mtb in 2D cultures at an MOI of 40 resulted in negligible infection ($n$ = 3).

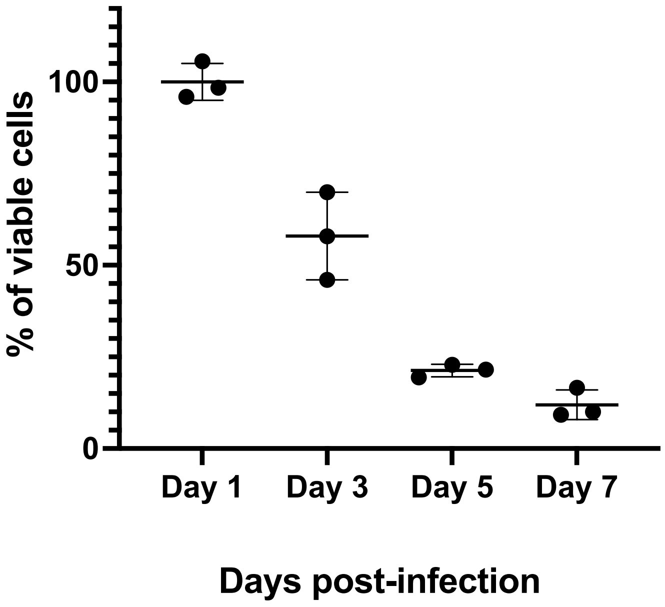


**Fig. S3 Viability of THP-1 macrophages infected with H37Rv in 2D cultures**. Percentage of viable cells at each time point was calculated as the percentage of live cells on Day 1.

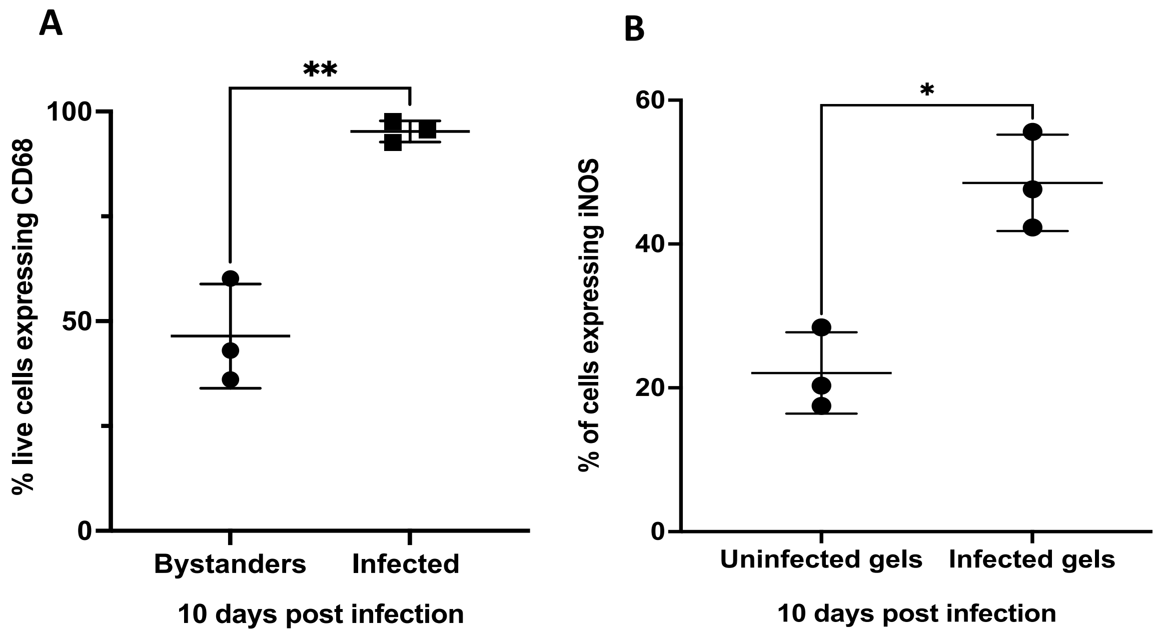


**Fig. S4 Infected cells differentiate into macrophages**. (**A**) Quantification of CD68 expression by bystanders and infected cells in collagen gels (*n* = 3). (**B**) Quantification of iNOS expression by THP-1 cells in uninfected and infected collagen gels (*n* = 3). Data in graphs represent the mean ± s.d., and *p* values were determined by two-tailed unpaired t-tests using GraphPad Prism Software. *p*-value < 0.05 was considered significant. ***p*=0.0026 for A, * *p*=0.0462 for B.

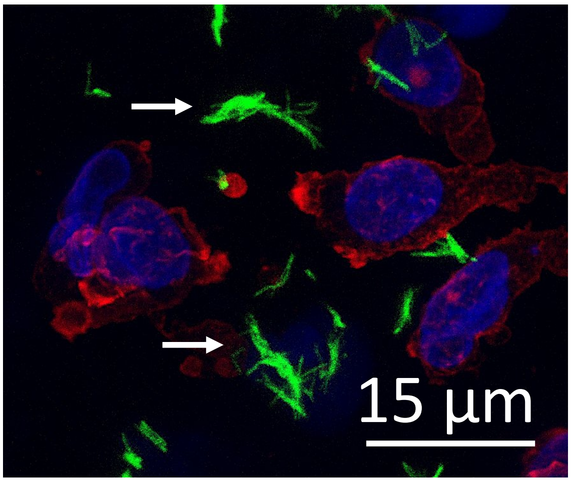

**Fig. S5 Representative image of extracellular cords (shown using arrows) 14 days post-infection.** DAPI (blue), H37Rv (Green), Phalloidin (Red).

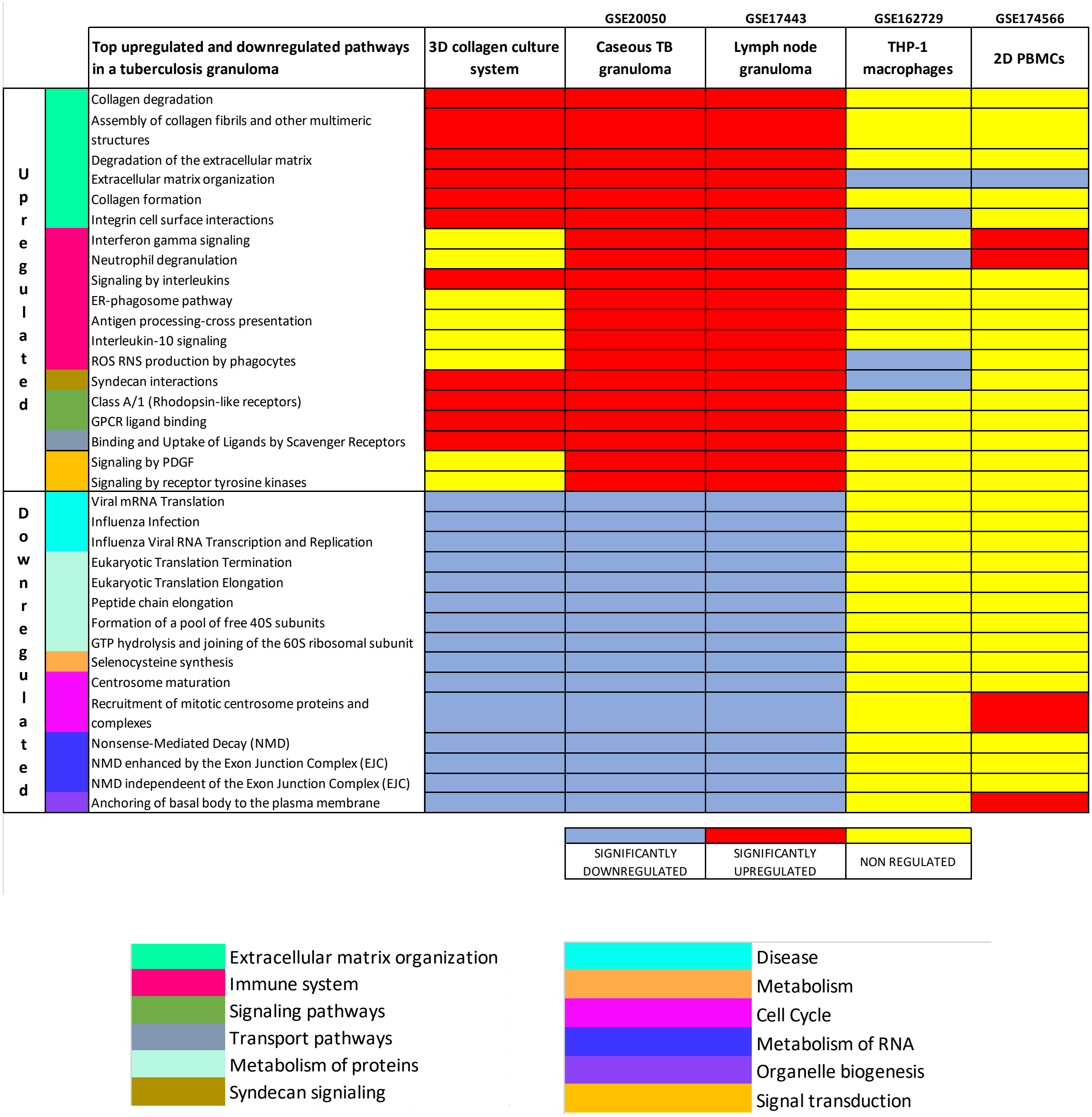


**Fig. S6 REACTOME pathway analysis of the mammalian cells from the infected collagen gels show maximum similarity with *in vivo* systems:** Pathways along with their gene ontology categories, in different culture conditions showing the 3D collagen culture system to be more representative of human *in vivo* infection (red: upregulated, blue: downregulated, yellow: non-regulated).

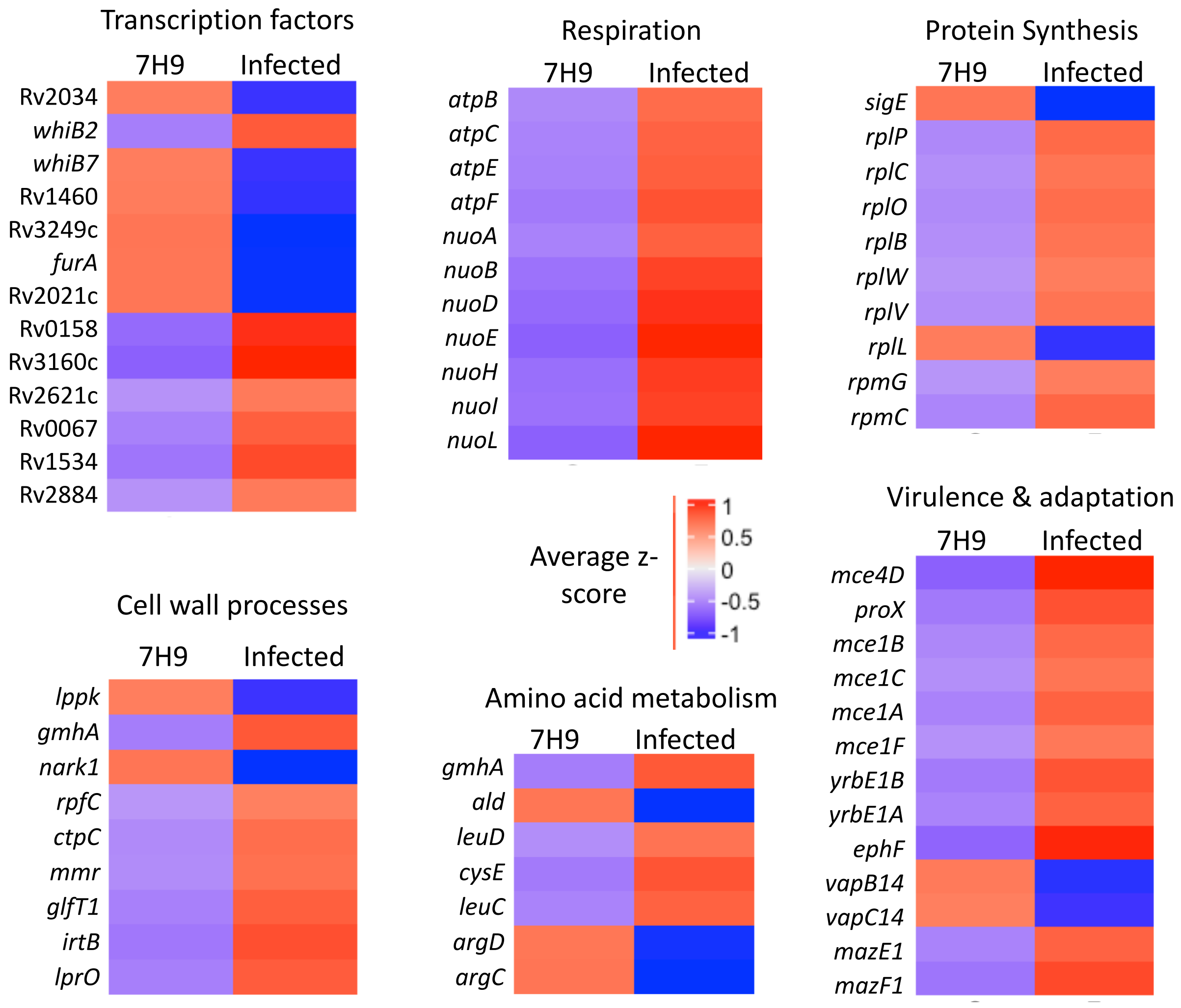


**Fig. S7.** Heat maps of differentially regulated Mtb belonging to various functional categories (source: Mycobrowser, École Polytechnique fédérale de Lausanne). Genes were considered differentially expressed on the basis of the false discovery rate (FDR) of ≤0.05 and absolute fold change of ≥1.)

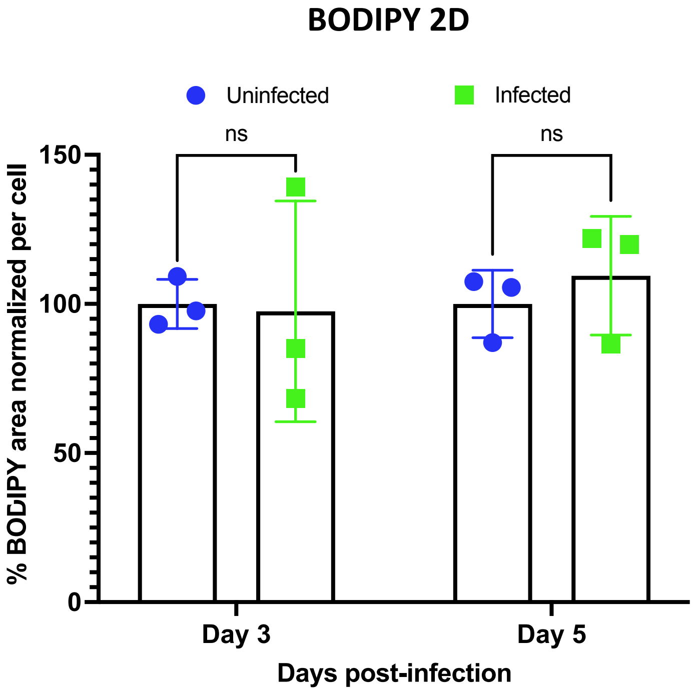


**Fig. S8 Infecting THP-1 macrophages in 2D with Mtb does not increase accumulation of lipid bodies in infected cells**. Plot showing total % BODIPY stained area normalized per cell at various time point after infection. Data in graph represents the mean ± s.d., and *p* values were determined by two-tailed unpaired t-test with Welch's correction using GraphPad Prism Software (ns: non-significant)

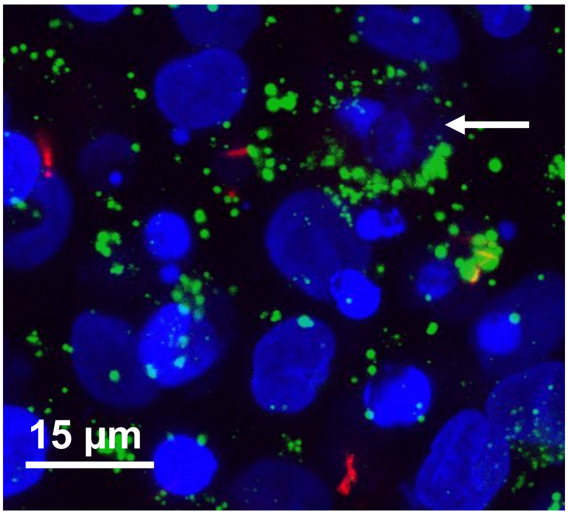

**Fig. S9 Mtb infection primes by-stander cells to acquire lipid bodies**: Representative images of by-standers THP-1 cells (shown by arrows) primed with lipid bodies (BODIPY-Green) in infected gels (DAPI- Blue, H37Rv – Red)

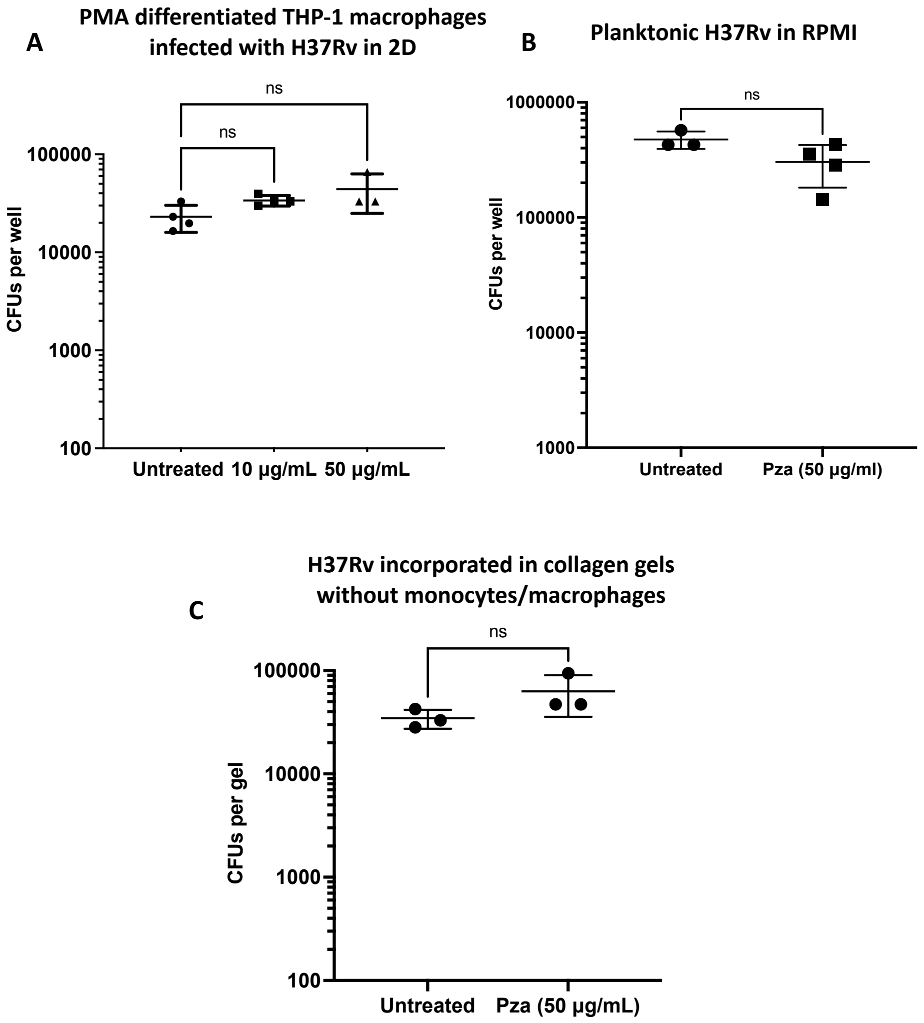


**Fig. S10 Pyrazinamide treatment is ineffective in reducing bacterial counts under various conditions.** CFU plots obtained after treating (**A**) PMA differentiated THP-1 cells infected with Mtb at 10 and 50 µg/mL for 3 days (*n* = 3 for 50 µg/mL and *n* = 4 for uninfected and 10 µg/mL), (**B**) planktonic bacteria suspended in RPMI (*n* = 3 for uninfected and *n* = 4 for infected) and (**C**) Mtb encapsulated in collagen gels without the THP-1 cells (*n* = 3). Data in graphs represent the mean ± s.d., and *p* values were determined by two-tailed unpaired t-test with Welch's correction using GraphPad Prism Software (ns: non-significant).

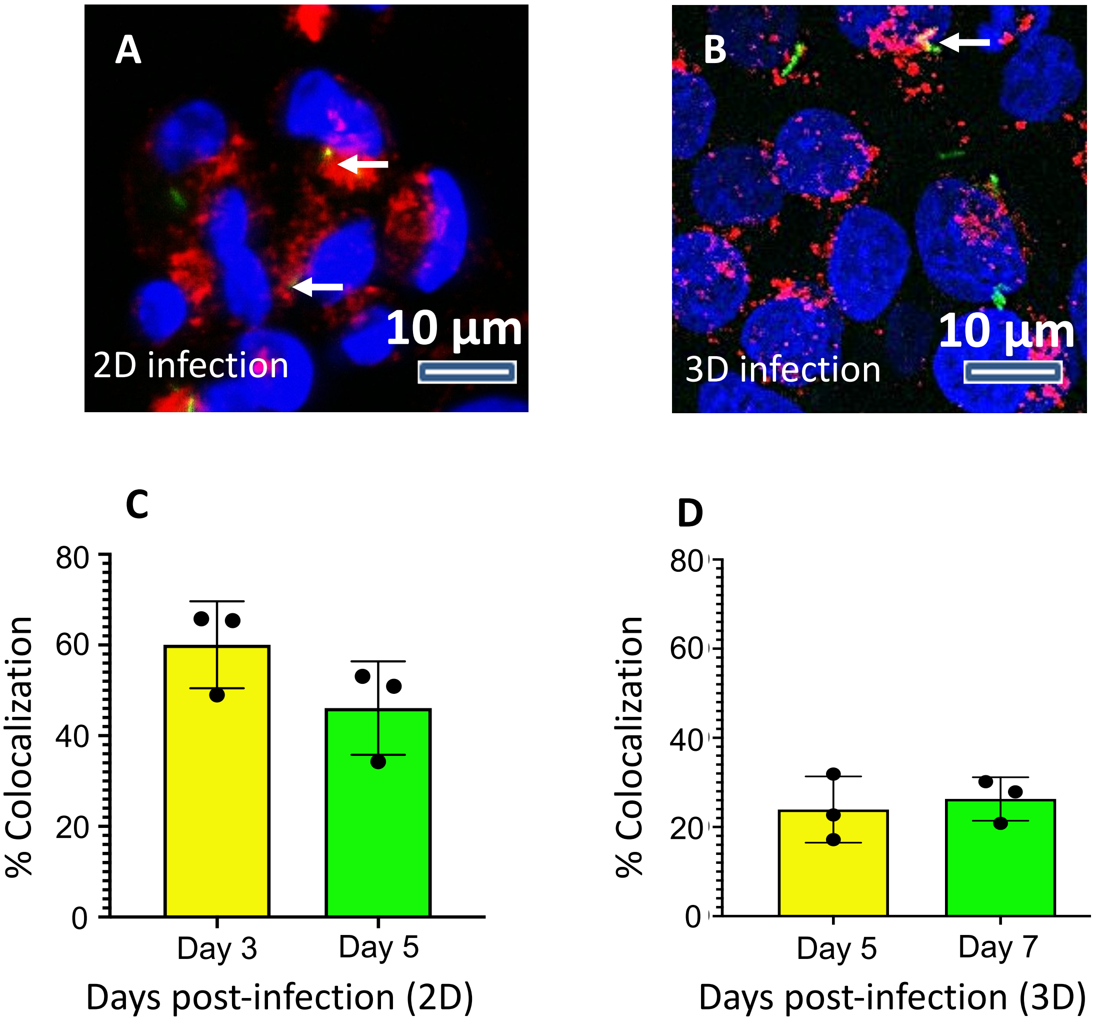


**Fig. S11 Colocalization of Mtb and late lysosomes (LAMP-1) did not significantly differ in 2D and 3D infection scenarios**. (**A**) & (**B**) Representative images of Mtb (green) infected THP-1 cells (nucleus: blue) stained with anti-LAMP-1 (red) in 2D and 3D respectively. Colocalized regions of Mtb and LAMP1 are in yellow (highlighted using arrows). (**C**) & (**D**) Quantification of Mtb and LAMP-1 colocalization in 2D and 3D infection respectively.

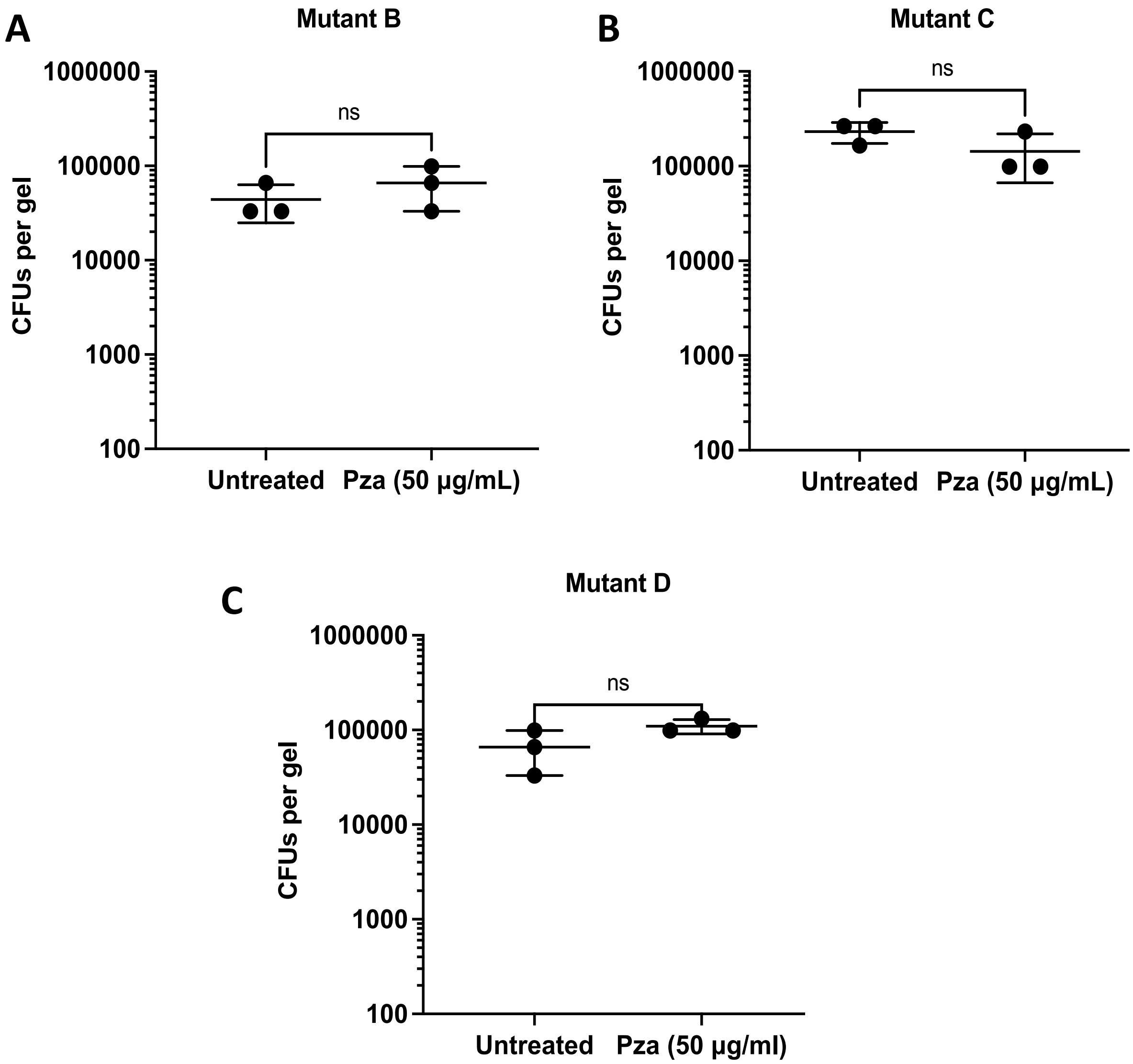


**Fig. S12 PZA was infective against Mtb H37Rv carrying mutations in the *panD* gene: (A-C)** Mtb (POA resistant strains) infected THP-1 cells with PZA at 50 µg/mL (ns: non-significant) ($n$ = 3), the treatment with PZA was started 4-days post-infection and was carried out for 6 days; hence the CFUs shown are 10 days post-infection. Data in graphs represent the mean ± s.d., and $p$ values were determined by two-tailed unpaired t-tests with Welch's correction using GraphPad Prism Software. $p$-value < 0.05 was considered significant. ns = non-significant. The mutations are listed in Supplementary Table 6.

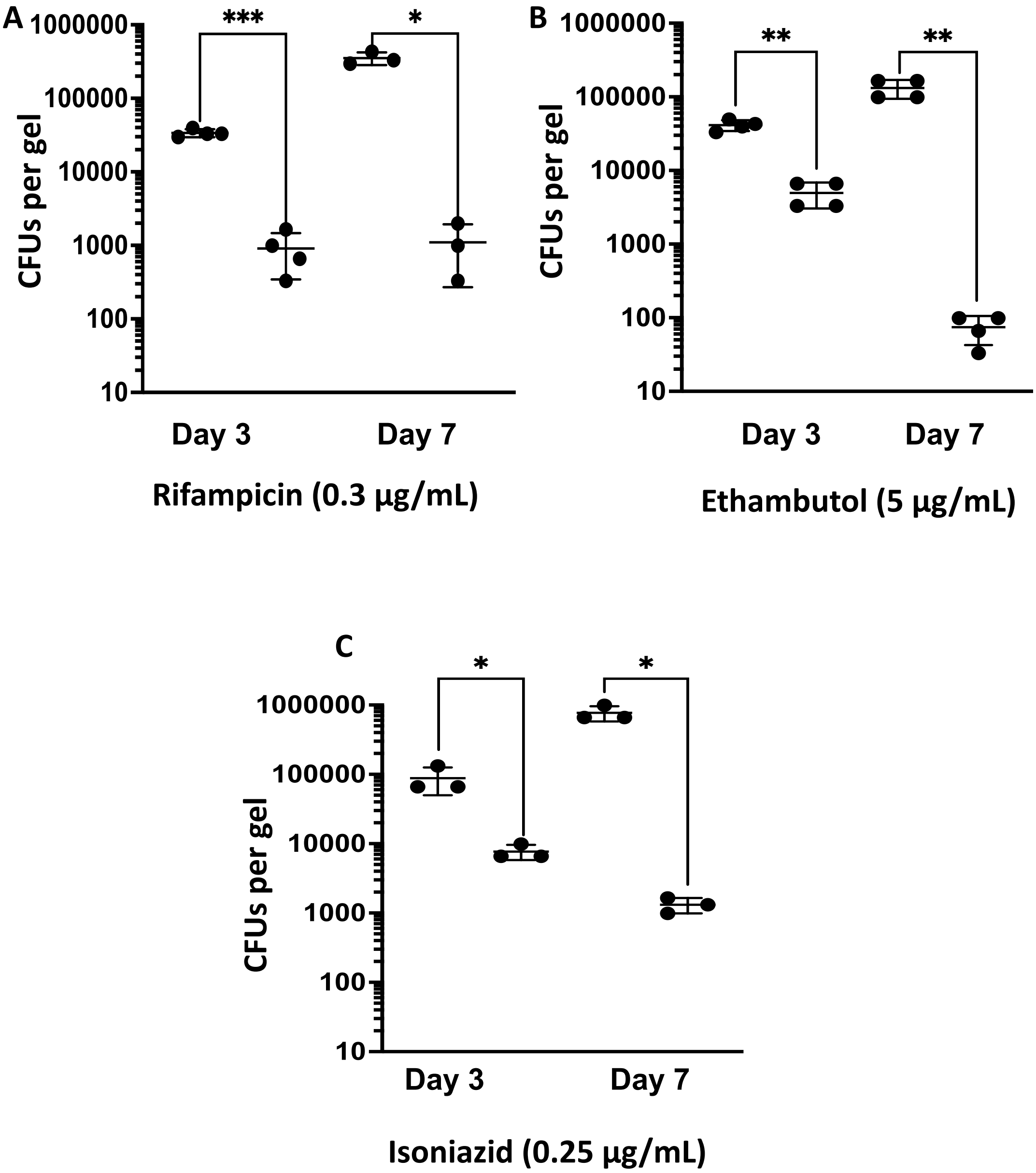


**Fig. S13 First line of tuberculosis drugs effectively reduces bacterial counts in collagen gels.** CFU plots obtained after treating with (**A**) Rifampicin ($n$ = 4 for Day 3 and $n$ = 3 for Day 7) (**B**) Ethambutol ($n$ = 4) and (C) Isoniazid ($n$ = 3). Data in graphs represent the mean ± s.d., and $p$ values were determined by two-tailed unpaired t-test with Welch's correction using GraphPad Prism Software. $p$-value < 0.05 was considered significant. For **A**: ***$p$=0.0005 for Day 3 and *$p$=0.0125 for Day 7, for **B**: **$p$=0.0011 for Day 3 and **$p$=0.0062 for Day 7, and **C**: $p$=0.023 for Day 3 and $p$=0.019 for Day 7.

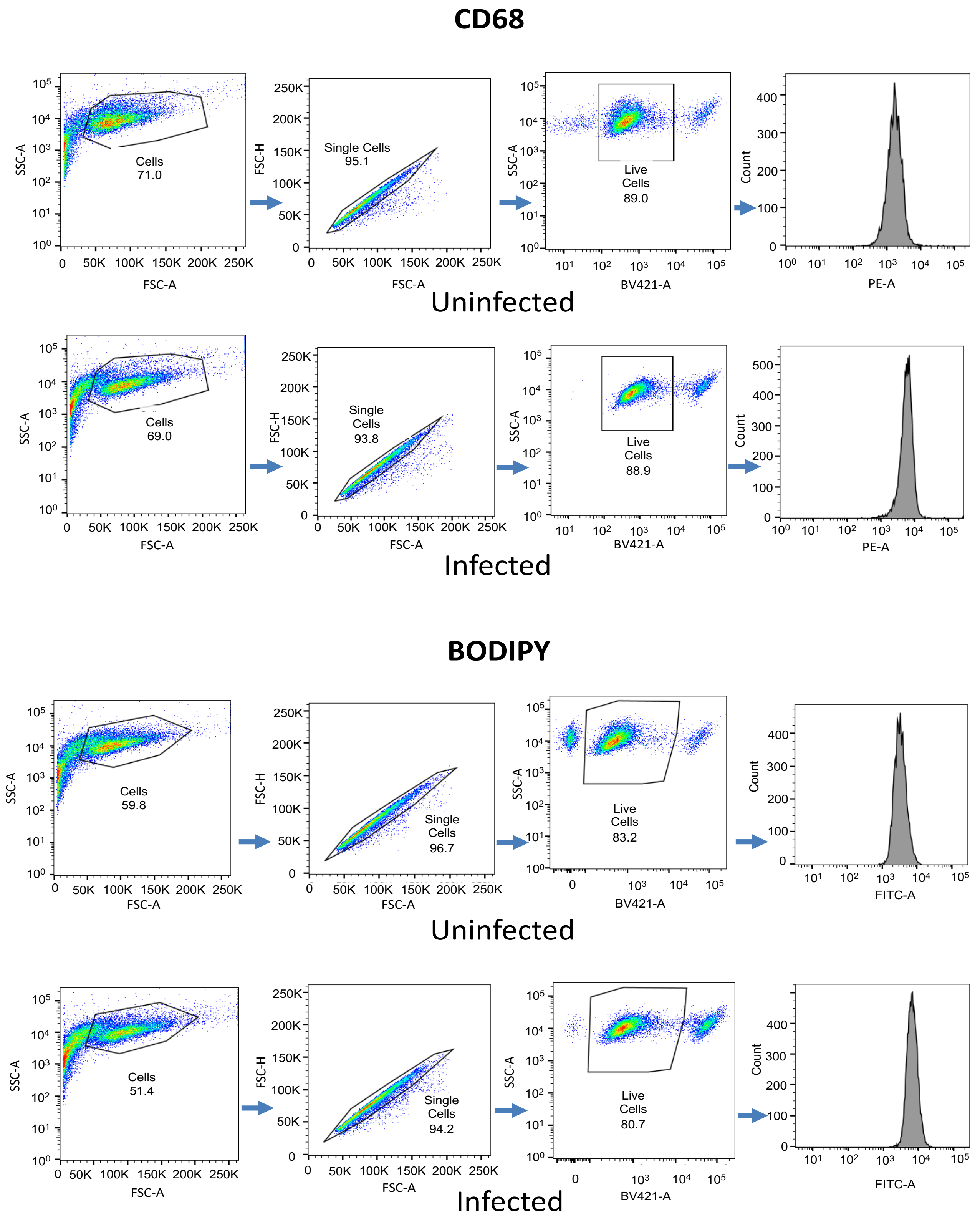


**Fig. S14 Flow cytometry gating strategy for CD68 and BODIPY staining.**

Flow cytometry gating strategy. Total THP-1 cells were first gated on a forward scatter (FS)/side scatter (SS) plot and then gated for single cells using forward scatter FSC-A/FSC-H. These were then further gated for live cells using BV421/SS. The live population was then gated for the infection based on the gating obtained for the corresponding uninfected population. Data were analyzed using FlowJo software, and population frequencies were expressed as percent of the parent population obtained in the live gating.

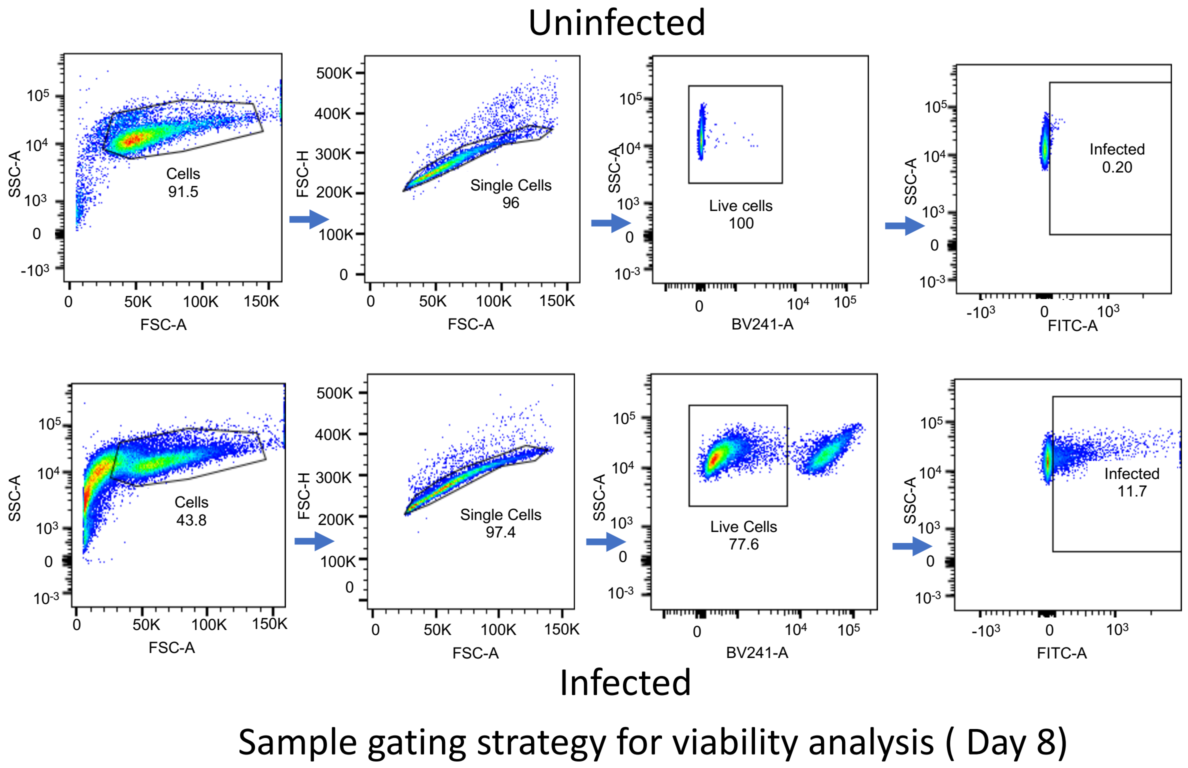


**Fig. S15 Flow cytometry gating strategy for viability assay.**

Flow cytometry gating strategy. Total THP-1 cells were first gated on a forward scatter (FS)/side scatter (SS) plot and then gated for single cells using forward scatter FSC-A/FSC-H. These were then further gated for live cells using BV421/SS. The live population was then gated for the infection based on the gating obtained for the corresponding uninfected population. Data were analyzed using FlowJo software, and population frequencies were expressed as percent of the parent population obtained in the single cells gate.
